# TEgment dissects transposable element reactivation upon CDK9 inhibition in acute myeloid leukemia

**DOI:** 10.64898/2026.09.16.752040

**Authors:** Dmitry Prokopov, Aisha Moussa, Hale Tunbak, Eve Leddy, Özgen Deniz

## Abstract

Cyclin-dependent kinase 9 (CDK9) inhibition represents a promising therapeutic strategy in cancers that disrupts RNA polymerase II pause release and collapses oncogenic transcripts. Here, we show that CDK9 inhibition also induces widespread transposable element (TE) activation across diverse cancer types. In acute myeloid leukemia (AML), CDK9 inhibition induces rapid epigenetic reprogramming and robust TE upregulation. To dissect this non-canonical transcriptional response, we developed TEgment, a TE-centric RNA-sequencing analysis pipeline that classifies TE expression into distinct structural modalities: standalone, readthrough, protein-coding gene-embedded (5′UTR, coding exons, and 3′UTRs), lncRNA-embedded, intron-retained, and TE-initiated/terminated transcription events. Applying TEgment to CDK9-inhibited AML revealed that TEs are predominantly expressed as readthrough, 3’UTR-embedded, and lncRNA-embedded transcripts rather than autonomous units, challenging prevailing assumptions about TE reactivation. TEgment provides a versatile framework for resolving TE transcriptional modalities at locus and structural levels, enabling precise interpretation of TE-derived transcripts in cancer and other contexts.

**Highlights:**

- CDK9 inhibition induces widespread TE expression across diverse cancer models
- AML cells show rapid, transient TE induction coupled to epigenetic reprogramming
- TEgment classifies TE-derived transcripts into structural modalities
- TEgment reveals predominant activation of TEs embedded within host transcripts upon CDK9 inhibition

## Introduction

Cancer cells frequently become dependent on the sustained high-level expression of oncogenic transcription factors to maintain proliferation, a phenomenon termed transcriptional addiction^1^. A critical checkpoint within the transcriptional machinery is promoter-proximal pausing, during which RNA polymerase II (Pol II) temporarily stalls shortly after initiating transcription. Release of Pol II into productive elongation is mediated by the positive transcription elongation factor b (P-TEFb) whose catalytic subunit cyclin-dependent kinase 9 (CDK9) phosphorylates the Pol II C-terminal domain as well as the negative elongation factors NELF and DSIF^2^. Through this mechanism, CDK9 enables productive elongation and is particularly essential for maintaining expression of short-lived oncogenic transcripts, including *MYC*, *MYB*, and *MCL1*^3^. Consequently, CDK9 inhibition or degradation rapidly disrupts these transcriptional programs, inducing apoptosis in diverse tumor models. Selective CDK9 inhibitors have demonstrated promising preclinical efficacy across both hematological and solid malignancies^4–7^. Among these, acute myeloid leukemia (AML) has emerged as a particularly compelling therapeutic context, because it is an aggressive disease characterized by transcriptional addiction, with a 5-year survival of approximately 30% in adults^8^.

Beyond causing transcriptional collapse at highly expressed genes, CDK9 inhibition has been shown to induce genome-wide epigenetic reprogramming, increasing chromatin accessibility^9^ and the derepression of epigenetically silenced tumor suppressor genes through recruitment of the chromatin remodeler SMARCA4/BRG1^10^. These global chromatin changes likely affect not only protein-coding genes but also non-coding and repetitive regions of the genome. In fact, a limited number of studies have reported that CDK9 inhibition can activate transposable elements (TEs)^9–11^, which comprise approximately 46% of the human genome^12^ and are normally silenced by epigenetic mechanisms. Furthermore, recent analyses of large-scale drug-perturbation screens have identified CDK9 inhibitors among the most potent inducers of TE expression across diverse cancer cell lines^13^. However, the mechanisms underlying TE reactivation remain poorly defined. It is unclear whether increased TE expression results from direct activation of TE promoters or is an indirect consequence of transcriptional readthrough and RNA processing defects. AML is a particularly important context in which to resolve this question. Its widespread epigenetic dysregulation and strong dependence on transcriptional control make CDK9-inhibited AML an ideal model for studying TE expression and transcript structure.

The technical complexity of quantifying TE expression from RNA-sequencing (RNA-seq) data presents a significant barrier to understanding TE biology in cancer. TE-derived signals can arise through several distinct mechanisms, including autonomous TE promoter activation, TE-initiated transcription, exonization, intron retention, and passive accumulation resulting from transcriptional readthrough; although these processes have different biological interpretations, they are often collapsed into a single measure of “TE expression”. Traditional approaches often overlook the remarkable heterogeneity of TE-derived sequences, conflating functional TE expression (e.g., autonomous, chimeric, exonized, or embedded within mature protein-coding and long non-coding RNAs (lncRNAs)) with passive intronic accumulation and transcriptional readthrough. Moreover, total RNA-seq captures both mature and immature transcripts but generates substantial intronic reads that require careful bioinformatic handling^14^. Specialized computational approaches are therefore necessary to contextualize TE signals within precise transcriptional landscapes and filter out spurious signals arising from incomplete RNA processing. Here, we present TEgment, a comprehensive RNA-seq analysis pipeline designed to classify TE expression events with high structural resolution.

Our pipeline categorizes TE transcripts into distinct modalities, including standalone transcription, readthrough events, lncRNA-embedded, untranslated region (UTR)-embedded, exon-embedded, exonized, intron-retained, and TE-initiated/terminated events, while removing unspliced pre-mRNA intronic artifacts. We applied TEgment to CDK9-inhibited AML as a model system to dissect the mechanisms driving TE reactivation and to place these events within a precise transcriptional and epigenetic context. Using this pipeline with the human telomere-to-telomere (T2T) genome assembly, we mapped these TE transcripts in CDK9-inhibited AML with unprecedented accuracy, revealing previously unannotated TE-derived transcript isoforms. Integrating TEgment with time-course RNA-seq and Cleavage Under Targets and Release Using Nuclease (CUT&RUN) revealed that early epigenetic reprogramming and downregulation of repressive chromatin regulators coincided with a rapid, transient induction of TEs upon CDK9 inhibition. Notably, TE reactivation was dominated by intragenic events, with TEs located within readthrough transcripts, embedded within 3′UTRs and lncRNAs emerging as the predominant transcriptional response to CDK9 inhibition.

Together, this approach provides a comprehensive framework for studying TE expression, which is broadly applicable across biological systems, and offers new insights into non-canonical transcriptional responses in AML.

## Results

### CDK9 inhibition induces widespread TE transcription across cancer cell lines

To determine whether TE activation represents a general transcriptional response to CDK9 inhibition, we first performed a systematic analysis of publicly available paired-end RNA-seq datasets (read lengths >100 bp). These datasets comprised 14 cell lines spanning 9 distinct solid and hematological cancer models treated with a range of CDK9 inhibitors and proteolysis targeting chimera (PROTAC) degraders across multiple time points (Table S1). Differential expression analysis using TElocal^15^, a locus-level TE quantification tool, revealed a pervasive TE dysregulation across most conditions, with a consistent bias toward upregulation (Figure 1A). In contrast, protein-coding genes exhibited a distinct pattern, with transcriptional downregulation frequently matching or exceeding upregulation across the same models (Figure S1A).

**Figure 1.**
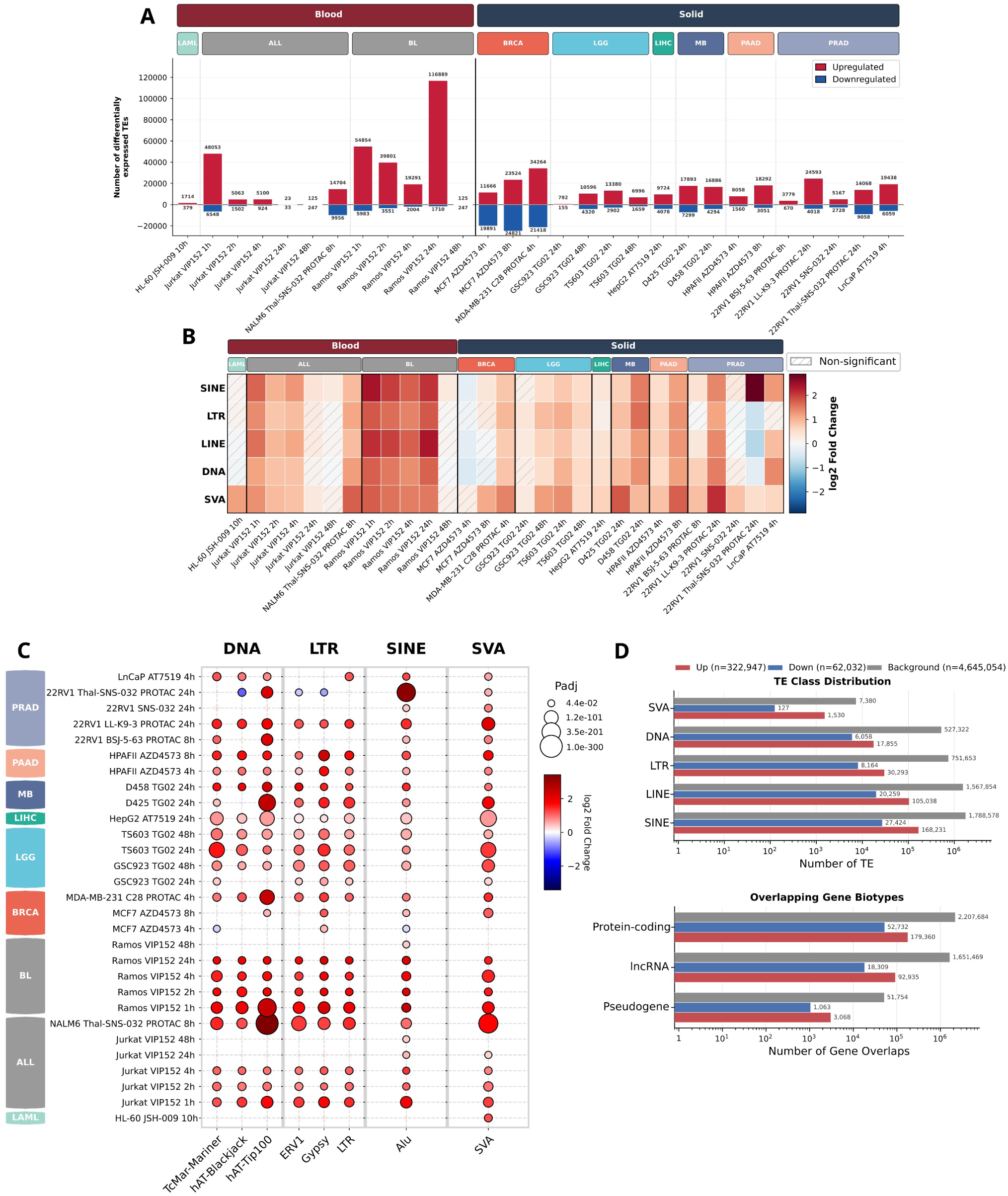
CDK9 inhibition drives widespread upregulation of TEs across diverse cancer models. (A) Numbers of differentially expressed TE loci following CDK9 inhibition. Data are grouped by tumor type (Blood vs. Solid) and cancer lineages: acute myeloid leukemia (LAML), acute lymphoblastic leukemia (ALL), Burkitt lymphoma (BL), breast invasive carcinoma (BRCA), lower-grade glioma (LGG), liver hepatocellular carcinoma (LIHC), medulloblastoma (MB), pancreatic adenocarcinoma (PAAD), and prostate adenocarcinoma (PRAD). Upregulated TEs are depicted in red and downregulated TEs in blue. Differential expression was determined using DESeq2 (adjusted p-value [padj] < 0.05, |log_2_FC| > 1.0). (B) Heatmap illustrating expression changes of TE classes across experiments. Color intensity corresponds to the mean log_2_ Fold Change and cells with hatching indicate non-significant expression changes (padj ≥ 0.05). (C) Differential expression of highly recurrent TE superfamilies across experiments. The color gradient indicates the log_2_ Fold Change (red indicating upregulation, blue indicating downregulation) and the size of each dot depicts the statistical significance (−log_10_(padj)). (D) Genomic characterization of consistently upregulated (red) and downregulated (blue) TE loci compared to the genomic background of all RepeatMasker-annotated TEs (gray). Top panel: Absolute frequency distribution of the deregulated TEs grouped by major TE class. Bottom panel: Distribution of deregulated TEs based on their overlap with genomic biotypes from GENCODE annotation. The x-axes are displayed on a logarithmic scale.

While TE dysregulation was observed in both solid and hematological tumors, the magnitude varied substantially across cellular context and treatment conditions. Next, we examined the recurrence of TE dysregulation across cell lines and treatment conditions, defining recurrent loci as those significantly changed in the same direction (|log_2_ FC| > 1.0) across multiple experiments. Across all datasets, 395,454 unique TE loci were differentially expressed; however, the majority were dysregulated in only a single experiment and were strongly biased toward upregulation (Figure S1B). Only a small subset (2.85%) exhibited recurrent activation across ≥5 experiments. Permuting significance calls within each experiment, matched for expression level, confirmed that this recurrence exceeds chance expectations (4.6-fold at ≥5 experiments, P = 1 × 10⁻⁴, Figure S1C). Protein-coding genes displayed a substantially higher degree of recurrent dysregulation across different conditions (Figure S1D), although the enrichment over their respective null distributions was comparable (4.2-fold, Figure S1E). To determine whether specific TE classes contributed to this response, we stratified differential expression by TE class. CDK9 inhibition resulted in a broad upregulation across all TE classes-SINEs, LTRs, LINEs, DNA transposons and SVAs (SINE-VNTR-Alus) (Figure 1B), indicating a generalized loss of transcriptional silencing rather than selective activation of a single TE type. Superfamily level analysis (Figure 1C), restricted to highly expressed elements (baseMean ≥ 500) with recurrent dysregulation in ≥ 11 experiments, confirmed that TE derepression involves upregulation of diverse superfamilies, ranging from evolutionarily ancient DNA transposons to young hominid-specific SVA elements. At the family level, applying stringent criteria with significance across ≥ 17 experiments, we identified 61 upregulated TE families, most dominated by LTR class, specifically the ERV1 superfamily (Figure S2A). Furthermore, SINE and DNA families exhibited consistent and high-magnitude upregulation across samples, while LINE and LTR families displayed more heterogeneous responses with downregulation in some conditions.

To further examine the genomic context of TE activation upon CDK9 inhibition, we intersected differentially expressed TE loci with genomic annotations. This analysis revealed that TE reactivation is significantly enriched within intronic regions, while depleted in intergenic, UTR, and exonic regions (Figure S2B). This intragenic distribution is predominantly associated with protein-coding host genes, followed by a substantial fraction mapping to lncRNAs (Figure 1D).

The specific accumulation of upregulated TEs within introns suggests that these elements are likely detected as part of unspliced pre-mRNAs or become exonized during splicing. Collectively, these analyses demonstrate that CDK9 inhibition triggers widespread yet context-dependent TE transcription across diverse cancer cell types, predominantly arising from intragenic loci and spanning multiple TE classes and families. However, this intragenic enrichment also exposes an interpretive limitation: standard locus-level quantification cannot distinguish bona fide TE transcription from unspliced pre-mRNA, exonized fragments, or readthrough inclusion, motivating the development of a dedicated structural classification framework.

### CDK9 is a critical dependency and prognostic determinant in AML

Given the widespread transcriptional dysregulation induced by CDK9 inhibition, we next asked whether CDK9 represents a critical transcriptional dependency in specific cancer types. AML is characterized by a strong dependence on transcriptional regulation, prompting us to investigate the expression and functional importance of CDK9 in this disease. Pan-cancer analysis of normalized The Cancer Genome Atlas (TCGA) expression data revealed that CDK9 is widely expressed across malignancies, with AML exhibiting elevated expression levels relative to most solid tumor cohorts (Figure 2A). Further comparison of uniformly normalized expression data from UCSC Xena validated that CDK9 expression is significantly upregulated in TCGA AML patient samples (n=173) compared to healthy bone marrow controls from the Genotype-Tissue Expression (GTEx) database (n=70; p<0.0001) (Figure 2B), indicating a cancer-specific overexpression of CDK9. Furthermore, Kaplan-Meier survival analysis demonstrated significantly shorter overall survival for patients in the CDK9-high group (median overall survival (OS): 12 months) compared to the CDK9-low group (median OS: 20 months), linking CDK9 overexpression to adverse clinical outcomes in AML (Figure 2C).

**Figure 2.**
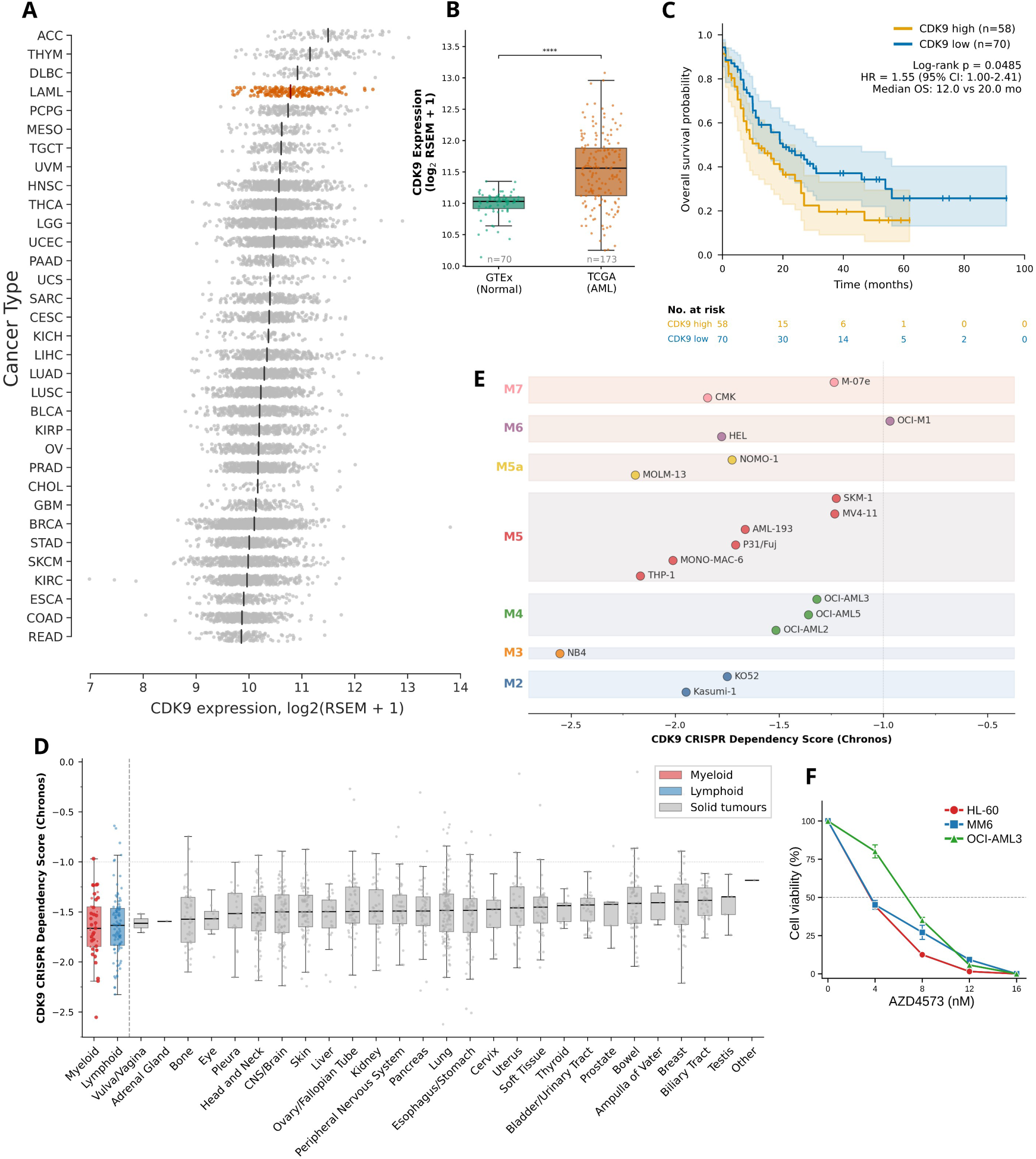
CDK9 is highly expressed and represents a critical vulnerability in AML. (A) Pan-cancer analysis of *CDK9* mRNA expression (log_2_(RSEM + 1)) using the TCGA PanCancer Atlas cohort. Acute Myeloid Leukemia (LAML) is highlighted in orange, others are shown in gray. Each dot represents an individual patient sample. Black line represents median value. (B) Comparison of *CDK9* expression between healthy bone marrow (GTEx, green, n = 70) and AML patient samples (TCGA, orange, n=173). Boxplots show the median and interquartile range (IQR), with individual patient samples overlaid as dots. ****p < 0.0001, determined by a two-sided Mann-Whitney U test. (C) Kaplan-Meier overall survival curve for the TCGA AML cohort, stratified by *CDK9* mRNA expression into *CDK9*-high (orange line, n = 58) and *CDK9*-low (blue line, n=70) groups. Shaded regions represent 95% confidence intervals. A table of the number of patients at risk is provided below the plot. (D) *CDK9* CRISPR dependency scores (Chronos) across cancer lineages using DepMap data. Boxplots depict median and IQR, with individual cell lines overlaid. Myeloid lineages are highlighted in red, lymphoid in blue, and solid tumors in gray. (E) Dot plot detailing *CDK9* CRISPR dependency Chronos scores for individual AML cell lines, categorized by FAB morphological classification (M2: blue, M3: orange, M4: green, M5: red, M5a: yellow, M6: purple, and M7: pink). The vertical dashed gray line indicates a Chronos score of -1.0. (F) Cell viability assay of three AML cell lines: HL-60 (red), MONO-MAC-6 (MM6; blue), and OCI-AML3 (green), treated with the CDK9 inhibitor AZD4573 (0–16 nM for 48 hours). The horizontal dashed line denotes 50% cell viability. Error bars represent standard deviation of three biological replicates.

We next investigated the functional dependency of cancer cells on CDK9 using data from the Cancer Dependency Map (DepMap) project, which demonstrated that CDK9 is highly essential across all cell lines. Notably, myeloid lineage cells exhibit some of the strongest dependencies on CDK9 across all cancer types (Figure 2D). Within AML, this essentiality is broadly conserved across diverse French-American-British (FAB) classifications (M2–M7), with Chronos scores, a CRISPR-based gene dependency metric modeling knockout effects on cell growth^16^, consistently falling below −1.0, establishing CDK9 as a critical survival requirement in these malignancies (Figure 2E). Consistent with these dependency data, treatment of three AML cell lines (OCI-AML3, HL-60, and MONO-MAC-6) with the highly selective CDK9 inhibitor AZD4573^17^ demonstrated high drug sensitivity, with complete cell death observed within 48 hours at low doses (16 nM, Figure 2F), further validating the critical dependency of AML cells on CDK9.

### CDK9 inhibition induces rapid transcriptional reprogramming

To interrogate transcriptional dynamics underlying CDK9 dependency in AML, we performed a time-course RNA-seq analysis in an AML cell line (MONO-MAC-6, FAB M5) treated with CDK9 inhibitor AZD4573 for 0, 4, 8, 24, and 48 hours. Likelihood ratio test (LRT) identified 10,301 significantly differentially expressed protein-coding genes (False Discovery Rate (FDR) < 0.05) across the time course (Figure 3A). Unsupervised k-means clustering (k=5) revealed transcriptional changes stratified into distinct temporal patterns, each associated with specific biological processes (Figure S3A).

**Figure 3.**
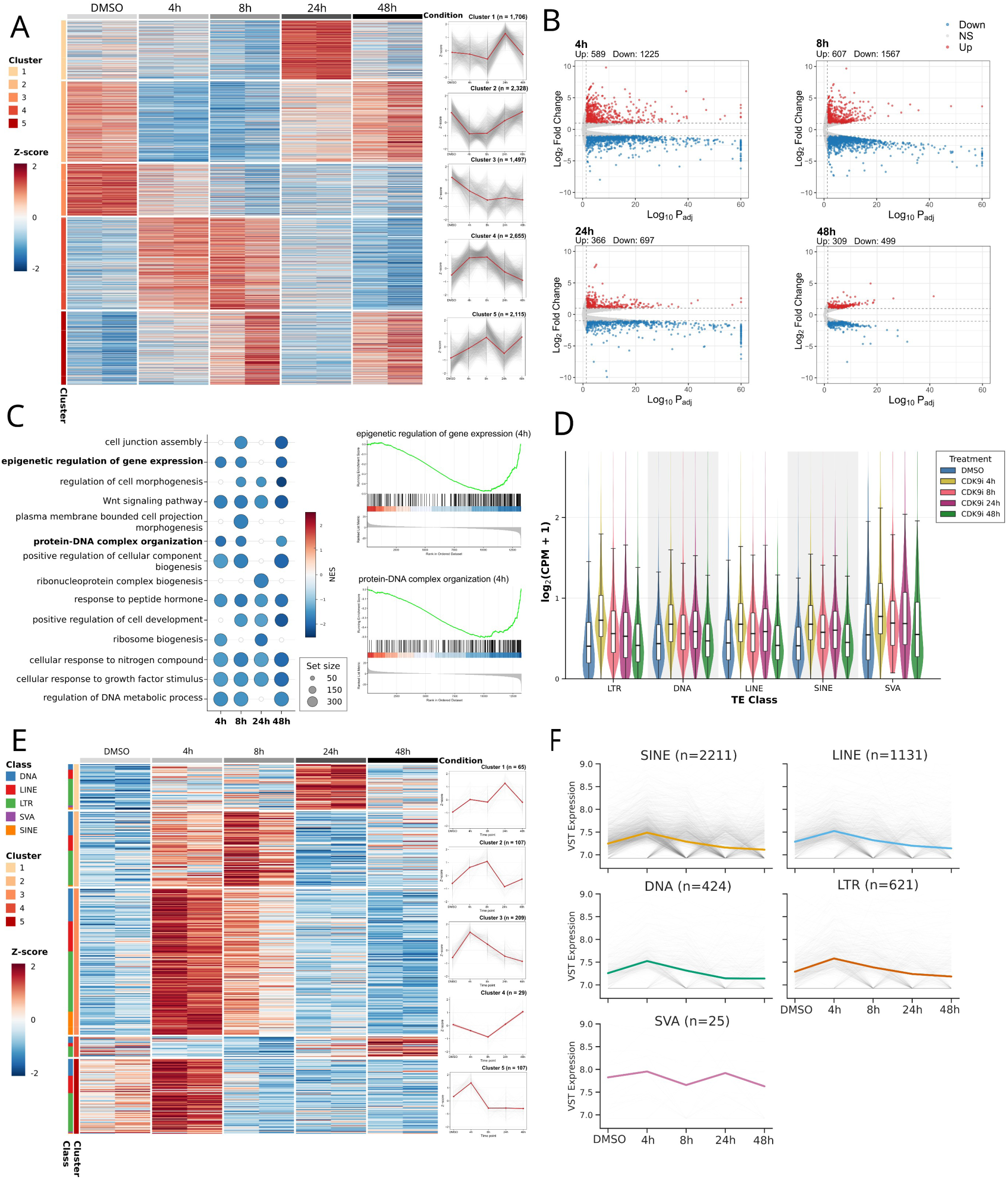
CDK9 inhibition induces transcriptional reprogramming and dynamic reactivation of TEs in AML. (A) Temporal expression profile of significantly differentially expressed protein-coding genes (LRT, FDR < 0.05) in MONO-MAC-6 cells treated with AZD4573. Heatmap shows z-scored, variance-stabilized expression across time points, clustered into 5 groups (colored bar). Right panels show average z-score trajectories per cluster, with the gray lines representing individual genes, red line indicating the cluster mean and the shaded area representing the standard error. (B) Pairwise differential gene expression relative to DMSO control. Significantly upregulated (Up, red dots) and downregulated (Down, blue dots) genes are defined by an absolute log_2_ fold change > 1.0 and adjusted p-value < 0.05. Non-significant (NS) genes are shown in gray. (C) Gene Set Enrichment Analysis (GSEA) results for significantly downregulated biological processes across the time course. Dot size corresponds to the number of genes in each set, and color intensity reflects the Normalized Enrichment Score (NES). Right panels show representative GSEA enrichment plots for “Epigenetic Regulation of Gene Expression” and “Protein-DNA Complex Organization” pathways at 4 hours. (D) Distribution of log_2_(CPM + 1) expression for expressed TE loci, categorized by TE class. Within each class, expression is shown across the time course: DMSO (blue), 4h (yellow), 8h (red), 24h (purple), and 48h (green). Box plots within violins indicate the median and IQR. (E) Expression dynamics of significantly differentially expressed TE families (LRT, FDR < 0.05) across the time course. Rows represent individual TE family, variance-stabilized and z-scored. TEs are grouped by k-means clustering (k=5). The colored bars on the left indicate the TE class and the assigned cluster. (F) Temporal expression trajectories (variance-stabilized expression) of individual significant TE loci, separated by TE class. Gray lines represent individual loci, while colored lines indicate the mean expression.

The largest cluster 4 (C4, n=2,655) demonstrated an early, transient induction of genes peaking at 4–8 hours with expression returning to baseline levels by 48 hours, including genes from HOX clusters and an enrichment of KRAB zinc finger (KZNF) genes (odds ratio = 1.43, FDR = 0.01; Table S2). Over-representation analysis (ORA) in this cluster revealed enrichment for regulation of immune and cytotoxic responses, suggesting acute stress-response activation upon CDK9 inhibition (Figure S3A). Other upregulated clusters displayed delayed activation, with C1 (n=1,706) peaking transiently at 24 hours and enriched for cell cycle and DNA damage checkpoints. C5 (n=2,115) peaked at 8 and 48 hours and was associated with mitochondrial respiration and intrinsic apoptotic signaling. Among repressed clusters, C3 (n=1,497) displayed progressive and sustained transcriptional downregulation consistent with Pol II pause release inhibition and was linked to Wnt and Hippo signaling pathways, while C2 (n=2,328) demonstrated an early transient repression at 4–8 hours followed by recovery at 48 hours and was defined by chromatin organization and nucleosome assembly (Figure S3A).

Pairwise differential expression analysis revealed that downregulated genes consistently outnumbered upregulated genes at all time points (Figure 3B), with maximal repression at 8 hours (1,567 downregulated versus 607 upregulated genes), before gradually attenuating at 24 and 48 hours post-treatment. Gene Set Enrichment Analysis (GSEA) demonstrated an early and sustained reduction in anabolic metabolism and chromatin maintenance programs, including regulation of DNA metabolic process, ribosome biogenesis, nucleosome assembly, protein-DNA complex organization, and epigenetic regulation of gene expression (Figure 3C and S3B). To dissect the suppression of chromatin pathway, we analyzed transcription of genes annotated to the “epigenetic regulation of gene expression” pathway across all timepoints. This revealed 157 epigenetic regulators, 150 of which were significantly dysregulated, with most showing acute downregulation at 4 and 8 hours (Figure S3C). This coordinated repression includes DNA methylation enzymes, H3K9 methyltransferases, Polycomb group genes, histone deacetylases and components of the human silencing hub (HUSH) complex that maintain silencing of repetitive elements. Overall, these data demonstrate that CDK9 inhibition induces widespread transcriptional reprogramming characterized by early and sustained suppression of chromatin maintenance pathways, potentially destabilizing epigenetic control across the genome.

### TE activation is early and transient following CDK9 inhibition

Coincident with the rapid loss of repressive chromatin machinery transcripts upon CDK9 inhibition, we observed widespread TE expression in MONO-MAC-6 cells. TElocal analysis revealed coordinated upregulation across all major TE classes (LTR, DNA, LINE, and SINE) after CDK9 inhibition, with the induction peaking at 4 hours followed by attenuation by 48 hours (Figure 3D). Similar early induction kinetics were observed in Jurkat (T-cell Acute Lymphoblastic Leukemia) and Ramos (Burkitt lymphoma) cells treated with a CDK9 inhibitor^11^ (Figure S4A), indicating that the rapid, transient pattern of TE activation represents a generalizable response to CDK9 inhibition across hematological malignancies. Family level expression analysis identified 485 TE families significantly dysregulated across the time course (LRT, FDR < 0.05, Figure 3E). Clustering stratified these TE families into early, mid and late response clusters (Table S3). Early response TE clusters 3 and 5 (TC3 and TC5) displayed peak upregulation at 4 hours, returning toward baseline by 24–48 hours. These clusters were enriched for primate-specific L1P families, including Homininae-derived L1PA2– 3 elements, as well as mammalian L1M subfamilies, such as the ancient L1MEd families (∼150 million years old)^18^. TC5 also contained the human-specific LTR5_Hs family and its internal region HERVK-int, demonstrating rapid activation of evolutionary young, full-length LTRs upon CDK9 inhibition (Figure S4B). Notably, TC3 showed the highest density of SINE families across clusters, dominated by AluS and AluY subfamilies and including the human-specific AluYb9, highlighting a robust early response of recent Alu elements. TE families at TC2 demonstrated a delayed peak at 8 hours before declining and were enriched for older LINE1s (with simian-origin L1PA8 as the youngest), a diverse set of LTR families (including epigenetic therapy-induced LTR12C^19,20^ and ancient MER41 elements), and two SINE families, revealing a mid-response cluster characterized by mixed-age LINE/LTR activation. A smaller subset of families in TC1 and TC4 exhibited sustained late induction at 24–48 hours. These clusters include both primate-specific L1PA6 (TC4), LTR10C and LTR7 families (TC1), as well as older LINE and DNA families such as L1MC3, L1MD2, MER1B and MER2 (TC1). Together, these patterns demonstrate that CDK9 inhibition triggers broad TE derepression across evolutionary ages and lineages rather than selective activation of specific classes.

At the locus level, we identified 4,412 significantly dysregulated TE elements across all time points using TElocal (Table S3). Locus level expression patterns recapitulated family level dynamics, with most loci showing transient induction at 4 and 8 hours followed by a return towards baseline by 24–48 hours (Figure 3F). Consistent with their genomic abundance, SINEs accounted for the largest fraction of responsive loci, dominated by AluSx family (495 loci). Notably, while most AluSx copies belonged to early-response clusters, individual AluSx loci were distributed across all five temporal clusters, with some exhibiting delayed or attenuated activation, indicating locus-specific regulation superimposed on shared sequence ancestry. LINE1 elements were also abundant, dominated by mammalian L1M elements (512 loci) with younger full-length LINE1s also present (Figure S4C) and showed predominantly early, transient activation. In contrast, LTR loci were fewer but followed comparable early- and mid-response patterns (Figure 3F). Collectively, these data demonstrate that CDK9 inhibition triggers rapid and transient TE derepression at thousands of loci, with temporal dynamics shaped by both subfamily identity and individual genomic context.

### TEgment reveals pervasive transcription of TEs embedded within host transcripts

While our initial pan-cancer analysis and time-course RNA-seq data in AML demonstrated a global burst of TE expression upon CDK9 inhibition, standard transcriptomic quantification methods present a significant technical barrier to interpreting these events. Traditional RNA-seq analytical approaches often conflate autonomous TE transcription with passive intronic signal or pervasive background transcription^21–23^, which is especially problematic in this context given the intragenic enrichment of upregulated TEs (Figure S2B), where many signals arise from host genes.

To place these events within their transcriptomic context, we developed TEgment, a comprehensive and modular short-read RNA-seq analysis pipeline designed to classify TE expression with high structural resolution (Figure 4A, Figure S5A). The workflow utilizes the complete T2T-CHM13v2.0 genome assembly as a reference. To overcome challenges of mapping highly repetitive sequences, TEgment provides flexible mapping strategies: users can restrict quantification strictly to uniquely mapped reads to identify high-confidence, unambiguous alignments, or optionally incorporate multi-mapping reads. For the latter, the pipeline employs expectation-maximization algorithms via TElocal to probabilistically resolve multi-mapping alignments, enabling accurate locus-level TE quantification.

**Figure 4.**
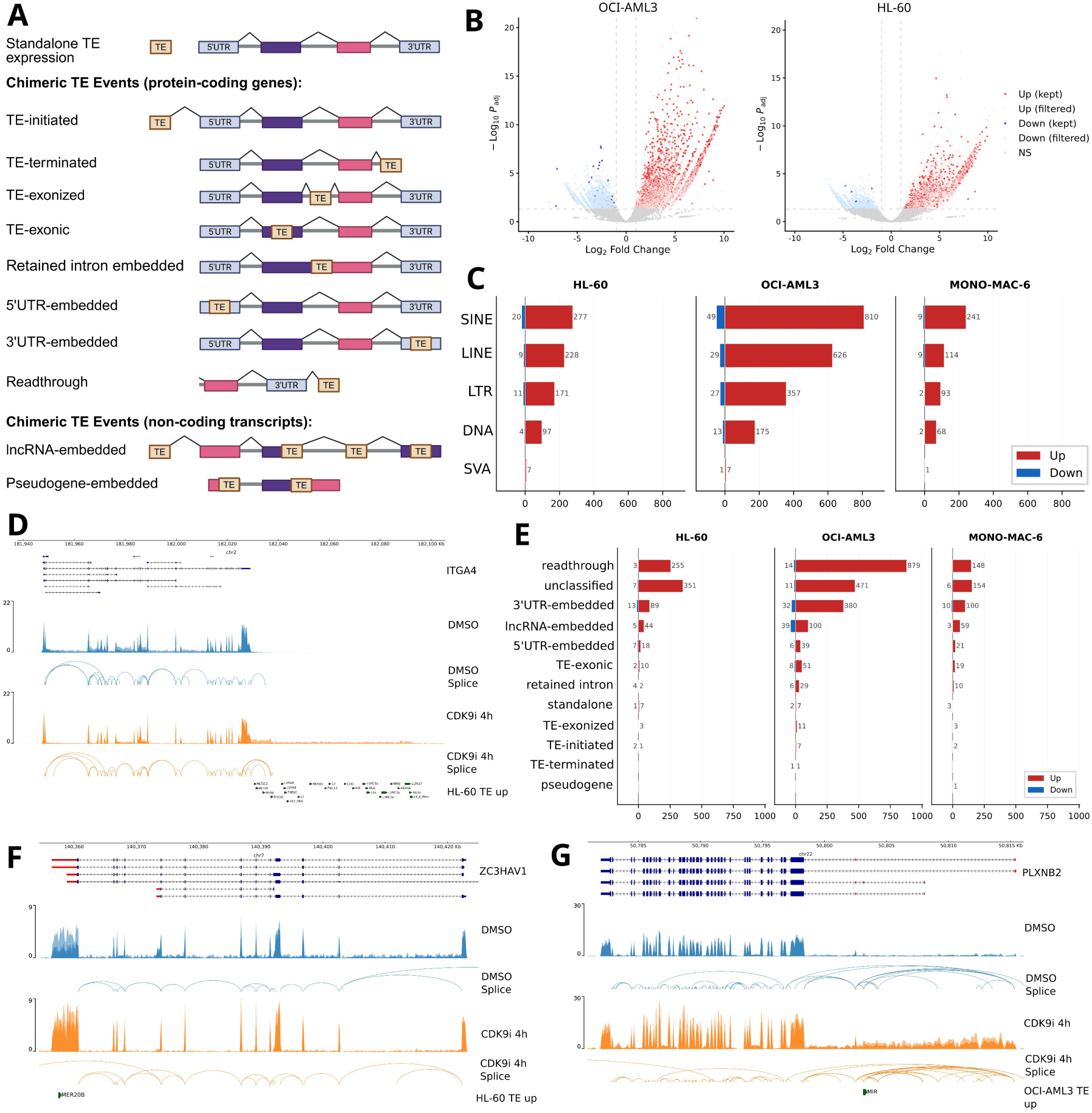
TEgment resolves the structural modalities of TE reactivation in AML following CDK9 inhibition. (A) Overview of the TEgment classification scheme, categorizing TE expression events into distinct structural modalities relative to host transcripts. (B) Locus level differential expression of TEs in OCI-AML3 and HL-60 cells following 4h CDK9 inhibition. Significantly dysregulated TEs retained after intersection with the assembled transcriptome are highlighted as solid red (upregulated) and blue (downregulated) dots (padj < 0.05, |log_2_ FC| > 1.0). Faded dots represent pervasive transcriptomic noise filtered out by the pipeline. (C) Absolute frequency distribution of significantly upregulated (red) and downregulated (blue) (padj < 0.05, |log_2_ FC| > 1.0). TEs stratified by major TE class across HL-60, OCI-AML3, and MONO-MAC-6 cell lines. (D) Representative genome browser track demonstrating transcriptional readthrough event in gene *ITGA4* upon CDK9 inhibition (CDK9i 4h, orange) compared to the DMSO control (blue). Tracks display normalized RNA-seq read coverage and splice junctions. Shaded regions indicate signal across three biological replicates. (E) Structural classifications of the significantly upregulated (red) and downregulated (blue) TEs across the three AML cell lines. (F, G) Genome browser tracks demonstrating distinct TE expression modalities upon CDK9 inhibition (CDK9i 4h, orange) compared to the DMSO control (blue). Tracks display normalized RNA-seq read coverage and splice junctions for (F) a 3’UTR-embedded TE in the *ZC3HAV1* gene, 3’ UTR marked in red, and (G) a 5’UTR-embedded TE in the *PLXNB2* gene, 5’ UTR marked in red. Annotated significantly dysregulated TE loci are indicated at the bottom of each panel. Shaded regions indicate signal across three biological replicates.

Following initial alignment, EASTR^24^ is applied to remove spuriously spliced alignments associated with repetitive elements. Subsequently, StringTie3^25^ performs reference-guided transcriptome assembly in a nascent-aware mode to accurately reconstruct expressed transcript isoforms. By intersecting locus-specific TE quantification with the newly assembled transcript models, TEgment categorizes TE expression events into distinct structural modalities. This distinguishes standalone transcription, TE-initiated, and TE-terminated events from TEs that are structurally embedded within host transcripts, including exonic, exonized, retained intronic, 5’UTR-embedded, 3’UTR-embedded, lncRNA-embedded, readthrough transcripts-embedded, and pseudogene-embedded events (Figure 4A, Figure S5B). TEs that do not fit any of these categories are annotated as “unclassified”. A primary advantage of TEgment is its dual capacity for structural detection and statistical quantification. By integrating structural annotation with robust differential expression analysis using DESeq2^26^, TEgment not only maps the architecture of TE-derived transcripts but also measures their significant expression changes across biological replicates.

We next applied TEgment to AZD4573-treated AML models to resolve the structural basis of TE-associated transcription following CDK9 inhibition. Given that TE upregulation peaked at 4 hours in MONO-MAC-6 cells (Figure 3D-F), we applied TEgment to two additional AML cell lines, HL-60 (FAB M2) and OCI-AML3 (FAB M4), treated with AZD4573 for 4 hours to determine whether this TE response is conserved across diverse AML models and to define the structural modalities of TE activation. At the global gene level, differential expression analysis identified a conserved response comprising 198 significantly upregulated and 298 downregulated genes shared across all three AML models upon treatment compared to Dimethyl sulfoxide (DMSO) control (Figure S5C). Both overlaps exceeded the permuted expectation (upregulated, 22-fold; downregulated, 6.7-fold; P = 1 × 10⁻⁴), supporting a conserved response (Figure S5G).

Prior to structural filtering, the initial locus level quantification identified a broad set of dysregulated TEs, heavily skewed by background intronic signal (HL-60: 1,498 upregulated, 5,780 downregulated; OCI-AML3: 3,620 up, 12,900 down; MONO-MAC-6: 2,143 up, 521 down). Intersecting these loci with the StringTie3-assembled transcriptome effectively filtered out this pervasive transcriptomic noise (Figure 4B, Figure S5D). This intersection narrowed the datasets, reducing the total number of reported dysregulated loci and revealing a consistent trend of TE upregulation across all three cell lines (HL-60: 780 up, 44 down; OCI-AML3: 1,975 up, 119 down; MONO-MAC-6: 517 up, 22 down) (Figure 4C). Despite this general trend, assessing the overlap of these loci demonstrated that TE dysregulation was highly cell-line specific, with only 55 loci consistently shared across all three cell lines (Figure S5E). This overlap still exceeded the permuted expectation 5.0-fold (P = 1 × 10⁻⁴), indicating that TE activation is largely cell-line specific (Figure S5G).

Stratification by major TE class showed upregulation across SINE, LINE, LTR, and DNA elements, with OCI-AML3 exhibiting the highest overall TE dysregulation. (Figure 4C). Among these classes, SINEs were the most frequently upregulated elements. At the family level, Alu (e.g., AluSx, AluJb subfamilies) and Mammalian-wide interspersed repeats (e.g., MIR, MIRb) were consistently enriched families in all three cell lines (Figure S5F).

We next applied the TEgment classification module to map the transcriptomic context of these dysregulated loci. This analysis revealed that the most prevalent mode of TE expression was transcriptional readthrough, where Pol II fails to recognize canonical transcription termination sites and continues transcribing beyond annotated gene boundaries, producing extended aberrant transcripts that incorporate downstream TEs (Figure 4D). This mechanism dominated across all three AML models, with OCI-AML3 showing 879 upregulated TE-readthrough loci at 4 hours post-treatment (Figure 4E). Importantly, library diagnostics confirmed low intergenic alignment rates (<9%) and high strandedness (>0.87) across all samples and conditions (Figure S6), excluding genomic DNA contamination as a source of the observed readthrough and intergenic TE signal. In contrast, application of TEgment to unperturbed, cell cycle-synchronised diploid RPE1 cells revealed only a minor fraction of TEs in readthrough transcripts, with TE expression instead dominated by standalone, lncRNA-embedded and 3′UTR-embedded modalities (Guiducci et al. 2026, submitted), indicating that readthrough-embedded TE expression is a specific consequence of CDK9 inhibition rather than a general feature of TE transcription. This enrichment for readthrough transcription is mechanistically consistent with CDK9’s role in coordinating Pol II processivity, where CDK9 inhibition impairs proper 3′ end processing and termination^27,28^, leading to accumulation of such aberrant transcripts. Following readthrough events, the next most common structural modalities involved TEs embedded within UTRs. In OCI-AML3 cells, we detected 380 loci featuring 3’UTR-embedded TEs (Figure 4F), while 5’UTR-embedded TEs encompassed 39 loci in OCI-AML3, 21 in MONO-MAC-6, and 18 in HL-60 cells (Figure 4G). This disproportion largely reflects the difference in UTR length: in GENCODE annotated 3′UTRs are substantially longer than 5′UTRs (median 640 versus 181 bp) and contain 2.6-fold more annotated TE loci (Figure S6B).

TEgment further enabled detection and quantification of more complex TE expression modalities, including lncRNA-embedded TEs and unannotated lncRNA isoforms. In some cases, lncRNAs were initiated from full-length LTR elements functioning as lncRNA genes (e.g., HERVE element, Figure 5A). Furthermore, TEgment also uniquely resolved diverse structural events beyond simple embedding, including TE-exonization events (e.g., AluSx within *LYST* gene generating a previously unannotated protein-coding isoform, Figure 5B), TE-initiated transcripts (e.g., MLT1E1A-driven *C1orf162*, Figure 5C), and TEs residing in previously unannotated intron retention events (e.g., AluSx1 within *ZNF589* gene, Figure 5D). Strikingly, contrary to prevailing assumptions about TE reactivation via dormant, autonomous promoters, standalone TE transcription was rare (e.g., AluY loci; Figure 5E), challenging the dominant paradigm of TE biology.

**Figure 5.**
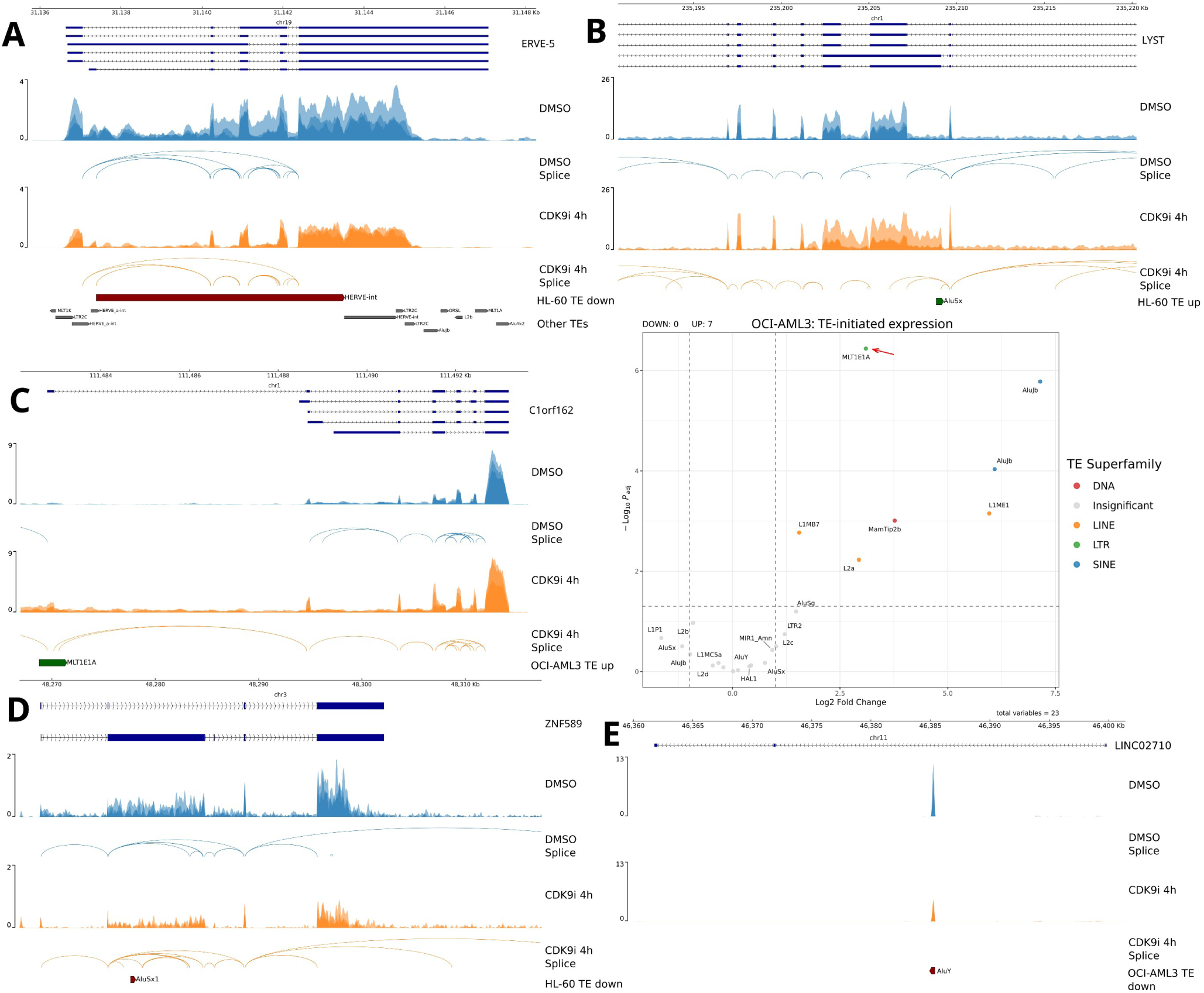
TEgment identifies complex structural modalities of TE expression. Genome browser tracks displaying normalized RNA-seq read coverage and splice junctions in AML cells treated with AZD4573 (CDK9i 4h, orange) versus DMSO control (blue). (A) The lncRNA transcript of the *ERVE-5* gene initiated from a full-length LTR element (HERVE). (B) TE-exonization event generating a novel protein-coding isoform (AluSx within *LYST*). (C) TE-initiated transcript demonstrating MLT1E1A-driven expression of the adjacent *C1orf162* gene (left). Volcano plot of TE-initiated events in OCI-AML3 cells following CDK9 inhibition (right). Significantly upregulated TEs (padj < 0.05, |log_2_FC| > 1.0) are coloured by superfamily (DNA, red; LINE, orange; LTR, green; SINE, blue), with non-significant elements shown in gray. Vertical and horizontal dashed lines denote the log_2_ fold change and significance thresholds, respectively. The MLT1E1A locus shown in the left panel is indicated (red arrow). (D) TE embedded within an intron retention event (AluSx1 within *ZNF589*). (E) Standalone TE transcription (AluY locus in the intron of *LINC02710* gene).

The enrichment of structurally embedded TEs and the minimal detection of standalone TE transcripts suggest that acute CDK9 inhibition does not primarily activate dormant, autonomous TE promoters. Instead, our data indicate that the disruption of transcriptional elongation, concurrent with the rapid downregulation of epigenetic silencers, results in the accumulation and inclusion of TEs that reside within actively transcribed host genes and lncRNAs. This suggests that, in AML, embedded TEs represent the predominant mode of TE activation upon CDK9 inhibition (Figure 4D). These examples illustrate TEgment’s capacity to resolve complex TE-gene architectures not captured by conventional quantification approaches. Together, they reveal a spectrum of TE-derived transcription, ranging from passive readthrough inclusion to novel TE-containing isoform generation and establish TEgment as a new framework for studying TE-host interactions across diverse biological contexts.

### CDK9 inhibition induces rapid loss of repressive chromatin marks

Having established that CDK9 inhibition triggers rapid downregulation of chromatin regulators (Figure S3C) and widespread TE activation (Figure 3E), we next sought to determine whether these transcriptional changes are associated with epigenetic reprogramming. Among the 298 genes commonly downregulated across all three AML cell lines at 4 hours post-treatment (Figure S5C), we identified core components of repressive chromatin complexes. These include core Polycomb repressive complex (PRC) components *EED* and *CBX2*, the histone deacetylase *SIRT1*, and the corepressor *LRIF1* (Table S2). This conserved disruption of silencing factors across AML models suggested that CDK9 inhibition might induce genome-wide epigenetic changes beyond its canonical effects on Pol II elongation. To test this, we performed Assay for Transposase-Accessible Chromatin using sequencing (ATAC-seq) and CUT&RUN in AML cells following 6 hours of AZD4573 treatment, focusing on the two cell lines exhibiting the most robust TE activation, OCI-AML3 and HL-60 (Figure 4C).

CDK9 functions canonically by phosphorylating the C-terminal domain of Pol II. Phosphorylation of Serine 5 (S5P) on the RPB1 subunit occurs shortly after transcription initiation when Pol II is at promoters, while phosphorylation of Serine 2 (S2P) marks elongating Pol II and increases towards the 3’ end region of transcribing genes^29,30^. To confirm the direct transcriptional effects of CDK9 inhibition, we performed CUT&RUN for Pol II-S5P, Pol II-S2P, and the active chromatin mark H3K27ac. As expected, CDK9 inhibition resulted in widespread loss of both Pol II-S5P and Pol II-S2P occupancy across transcription start sites (TSSs) and gene bodies in both cell lines (Figure 6A). However, the extent of Pol II phosphorylation loss differed between the two cell lines: while HL-60 cells exhibited near-complete loss of Pol II-S5P and S2P signals, OCI-AML3 cells showed partial retention of Pol II phosphorylation, suggesting differential sensitivity to AZD4573. Genome-wide analysis confirmed this pattern, showing reduction of Pol II-S5P, Pol II-S2P, and H3K27ac signals, which was stronger in HL-60 cells (26,454 loci for Pol II-S5P) compared to OCI-AML3 cells (15,708 loci for Pol II-S5P) (Figure 6B, Figure S7A-C). Pol II-S5P signal loss was relatively evenly distributed between TSS and non-TSS regions, suggesting that CDK9 inhibition also affects non-genic Pol II targets, including lncRNAs and TEs (Figure S7C). Notably, 71% of downregulated genes in HL-60 lost Pol II-S5P signal upon CDK9 inhibition (CDK9i), demonstrating that the most transcriptional repression is a direct consequence of impaired Pol II pause release. On the other hand, only 40% downregulated genes showed Pol II-S5P loss in OCI-AML3 (Figure S7D), suggesting that transcriptional repression involves both direct effects of impaired pause release and indirect consequences of disrupted transcriptional networks in this cell line. Nevertheless, among commonly affected genes was the known CDK9-dependent oncogenic transcript *MYB*; key epigenetic regulators, including *EED* (core Polycomb Repressive Complex 2 component), *SIRT1* (NAD-dependent histone deacetylase), and *SMARCAD1* (SWI/SNF-related chromatin remodeler), where S5P, S2P, and H3K27ac signals were substantially diminished at 6 hours post-treatment in both HL-60 and OCI-AML3 (Figure 6C, S8A-B).

**Figure 6.**
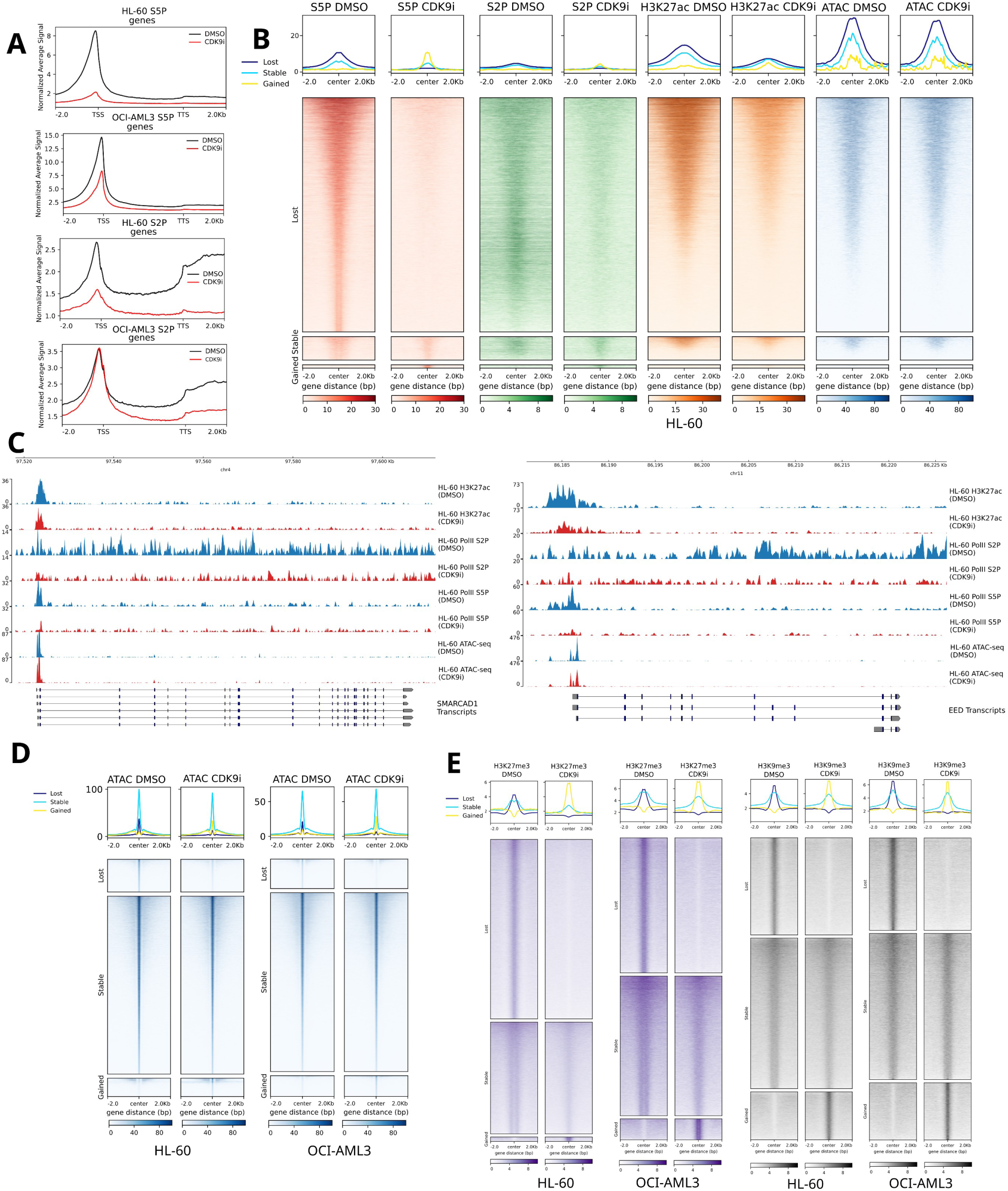
Acute CDK9 inhibition induces genome-wide epigenetic reprogramming. (A) Normalized continuous coverage of Pol II-S5P, Pol II-S2P, H3K27ac, and ATAC-seq across genic regions (± 2 kb flanking regions) in AML cells. Profiles compare DMSO control (black lines) to 6h AZD4573 treatment (CDK9i, red lines). (B) Heatmap showing normalized and IgG background-subtracted CUT&RUN and ATAC-seq signal intensities centered on S5P consensus peak regions (± 2 kb). Regions are stratified into Lost (log_2_ fold change ≤ -1), Stable, and Gained (log_2_ fold change ≥ 1) subsets upon 6h CDK9 inhibition based on Pol II S5P signal. (C) Genome browser tracks of CUT&RUN and ATAC-seq signal at the *SMARCAD1* (left panel) and EED (right panel) gene loci in HL-60. Tracks compare DMSO control (blue) and 6h CDK9i treatment (red) for H3K27ac, Pol II-S2P, Pol II-S5P, and ATAC-seq. (D) ATAC-seq signal intensity comparing DMSO to 6h CDK9i treatment across consensus peak centers (± 2 kb) across Lost, Stable, and Gained categories. (E) Genome-wide signal distribution of repressive marks H3K27me3 (purple) and H3K9me3 (gray) at peak centers (± 2 kb). Target regions are partitioned by response to CDK9i (Lost, Stable, Gained).

To test whether acute CDK9 inhibition affects chromatin structure, we performed ATAC-seq at 6 hours post-treatment. Surprisingly, global chromatin accessibility remained mostly stable, with no substantial change in overall ATAC signal in both HL-60 and OCI-AML3 cells (Figure 6D). Genome-wide comparison of ATAC peaks identified 11,724 and 21,081 regions with increased accessibility and 20,935 and 30,146 regions with decreased accessibility in HL-60 and OCI-AML3 cells, however these changes were comparatively weak, with the vast majority (113,546 and 182,164) of regions remaining unchanged. Notably, regions with Pol II-S5P loss upon CDK9 inhibition showed no significant change in chromatin accessibility (Figure 6B), overall indicating that chromatin compaction state is largely unaffected during the acute response. Although chromatin accessibility was largely unaffected, CUT&RUN profiling revealed substantial genome-wide loss of both H3K27me3 and H3K9me3 within 6 hours of CDK9 inhibition (Figure 6E). H3K27me3 depletion was more pronounced than H3K9me3 loss, aligning with the observed downregulation of multiple PRC components. Furthermore, this loss was more evident in HL-60 cells and extended to TE-enriched genomic regions (Figure S8C). On the other hand, H3K9me3 depletion was evident in both cell lines and predominant within TEs, with 55% and 58% of lost regions located within TEs in HL-60 and OCI-AML3 cells (Figure S8C).

Together, these epigenetic profiling experiments reveal that acute CDK9 inhibition triggers rapid, genome-wide loss of H3K27me3 and H3K9me3 without major chromatin accessibility changes. This early epigenetic reprogramming at TE-enriched regions coincides with the widespread but transient TE activation observed in CDK9-inhibited AML cells.

## Discussion

TEs constitute nearly half of the human genome yet remain among the most poorly characterized features of transcriptomes. While TE activation has been reported across diverse biological contexts, including development^31,32^, aging^33,34^, inflammation^35,36^, and cancer^37,38^, the technical challenges of accurately quantifying TE expression have limited our understanding of their functional roles. Importantly, RNA-seq signals from TE loci can arise from distinct transcript architectures, rather than a single molecular process and conventional TE quantification approaches collapse these structurally distinct events into a single measure of “TE expression”.

In this study, we address this gap by developing TEgment, a comprehensive computational pipeline that resolves TE expression at both locus and structural levels, distinguishing biologically meaningful TE activity from pervasive background noise. Applying TEgment to CDK9 inhibition in AML, we demonstrate that acute transcriptional perturbation triggers widespread TE activation, driven predominantly by TEs embedded within host transcripts.

### CDK9 inhibition as a model system for TE biology

CDK9 has emerged as a promising therapeutic target in oncology, with multiple inhibitors in clinical trials across hematological malignancies and solid tumors^39,40^. CDK9 inhibition has effects that extend well beyond canonical transcriptional repression: perturbing RNA processing, interfering with coordination of spliceosome recruitment, promoting readthrough transcription and inducing genome-wide chromatin remodeling^9^. A limited number of studies have reported TE activation following CDK9 inhibition^9–11^, yet the mechanisms and the extent underlying this response and its functional significance remain undefined.

Our systematic analysis of 14 cancer cell lines across 9 tumor types reveals that TE activation is a pervasive, reproducible response to CDK9 inhibition, affecting all major TE classes: LINEs, SINEs, LTRs, SVA and DNA transposons. It is the breadth and extent of transcriptional disruption that make CDK9 inhibition an ideal, but technically challenging, system for studying transcriptional dynamics of TEs. While CDK9 inhibition leads to a robust and reproducible TE signal, it also produces the full spectrum of transcriptional artifacts that confound conventional TE quantification approaches. Disentangling these contributions requires structural resolution of TE-derived transcripts, establishing CDK9 inhibition as both a compelling biological context and an ideal stress test for developing TEgment.

### Epigenetic reprogramming and transcriptional readthrough coincide with TE activation

Our integrated time-course RNA-seq and chromatin profiling experiments indicate that TE reactivation upon CDK9 inhibition is associated with two mechanistically distinct but temporally parallel processes: disruption of co-transcriptional RNA processing and rapid loss of repressive chromatin marks at TE-containing loci. The first mechanism is a direct consequence of CDK9’s canonical function. CDK9 coordinates Pol II elongation, 3′ end processing, and transcription termination through phosphorylation of the RNA processing machinery^28^. CDK9 inhibition therefore impairs transcript termination, causing Pol II to read through annotated gene boundaries and incorporate downstream TE sequences into aberrant transcripts. TEgment identified transcriptional readthrough as the largest category of TE activation across all three CDK9-inhibited AML models. This context-dependence underscores why structural classification is essential, as the same TE locus can be activated by entirely different mechanisms depending on the biological perturbation.

The second mechanism involves chromatin level changes and is less directly predicted from CDK9’s canonical role. Within 4–8 hours of CDK9 inhibition, we observe coordinated transcriptional downregulation of epigenetic silencing machinery, including core components of TE-silencing pathways: DNA methyltransferases (*DNMT1*, *DNMT3A*), chromatin remodeler (*SMARCAD1*), H3K9 methyltransferases (*SETDB1*, *SUV39H1*, *SETDB2*), Polycomb group genes (*EED*, *EZH1*, *CBX2/4/8*), and both the core components and interacting partners of the HUSH complex (*TASOR*, *MPHOSPH8*, *MORC2*). Coincident with this loss of silencing factors, we observed genome-wide redistribution of repressive histone marks, with substantial depletion of H3K27me3 and H3K9me3 at non-TSS regions, including TE loci, which may facilitate activation of TE-containing loci, including readthrough-derived transcripts.

Notably, while previous studies reported increased chromatin accessibility following CDK9 inhibition at 24–96 hours post-treatment^9,10^, our ATAC-seq analysis at 6 hours revealed stable global accessibility. This temporal difference suggests a biphasic chromatin response where early loss of repressive marks (detected by CUT&RUN) precedes subsequent chromatin decompaction (detected by ATAC-seq). Notably, TE upregulation is predominantly transient, peaking within 4–8 hours before attenuating by 24–48 hours, suggesting that active re-silencing mechanisms restore chromatin control even under sustained CDK9 inhibition.

Our CUT&RUN experiments also confirmed the expected acute effects of CDK9 inhibition on transcriptional machinery: loss of Pol II Ser2/Ser5 phosphorylation and H3K27ac at most downregulated genes, consistent with impaired pause release and transcriptional collapse at canonical genes. Notably, while loss of Pol II-S5P/S2P, as well as repressive H3K9me3 and H3K27me3 marks was more predominant in HL-60 cells, OCI-AML3 cells demonstrated more robust TE activation, highlighting both cell-type specific sensitivity and response to CDK9 inhibition in different AML models. This indicates that the chromatin changes captured by our CUT&RUN profiling are unlikely to fully account for the magnitude of TE-associated transcriptional dysregulation in OCI-AML3 cells, and that additional epigenetic mechanisms, potentially changes in DNA methylation, and other epigenetic pathways not examined in this study are likely to contribute to TE activation following CDK9 inhibition. Together, these observations argue that CDK9 inhibition elicits pleiotropic, cell-context-dependent epigenetic effects, and that TE reactivation is unlikely to be driven exclusively by H3K9me3 and H3K27me3 loss.

We also observed robust upregulation of over 100 KZNF genes within 4–8 hours of CDK9 inhibition. KZNFs are the largest family of transcriptional repressors in mammals and play critical roles in TE silencing^41–43^. Their induction might reflect a compensatory transcriptional response to restore silencing capacity as repressive chromatin collapses in order to prevent even more extensive TE activation. Alternatively, given that many KZNFs themselves contain TEs within their regulatory regions, their activation may be a direct consequence of local chromatin derepression. This feedback loop between TE activation and KZNF induction warrants further investigation.

### TEgment reveals that TE expression is largely explained by intragenic host-transcript architectures

A central challenge in TE biology is differentiating bona fide TE transcription from technical artifacts. Standard quantification approaches treat TE loci as independent features, collapsing reads into locus- or family-level counts without reference to the surrounding transcriptional architecture. Therefore, reads arising from unspliced pre-mRNA^23^, exonized TE fragments^22^, intron retention^44^, or transcriptional readthrough^21^ become indistinguishable from reads produced by TE-autonomous promoters. This ambiguity has hindered efforts to assign biological meaning to TE expression and has led to widespread overestimation of TE reactivation in published studies, many of which likely reflect host-transcript dysregulation rather than TE-autonomous activity.

TEgment was designed around a simple interpretive principle: every TE expression event must be evaluated in the context of the transcript that contains it. By coupling locus-resolved TE quantification to sample-specific reference-guided transcriptome assembly, the pipeline classifies each dysregulated locus not as an isolated count but as a structurally defined event: standalone, TE-initiated, TE-terminated, exonized, or embedded within a specific host-transcript context (5′UTR, 3′UTR, coding exon, retained intron, lncRNA, or readthrough extension). Notably, by using the recent T2T-CHM13v2.0 genome assembly and incorporating EASTR filtering to remove spuriously spliced alignments, TEgment maps TE-derived transcripts and distinguishes genuine expression of TEs from artefactual signal, arising from mis-mapped, spuriously spliced alignments.

Importantly, although TEgment filters transcript structures arising from incompletely processed or spuriously aligned transcripts to improve structural annotation of TE-derived transcriptional events, these RNA species should not necessarily be regarded as biologically irrelevant. CDK9 inhibition profoundly perturbs RNA polymerase II elongation and RNA processing, and accumulation of incompletely processed transcripts may itself represent a biologically meaningful consequence of this perturbation. TEgment was designed to distinguish these events from structurally resolved TE-associated transcripts, rather than to imply that the former lack biological significance.

Application of TEgment to three CDK9-inhibited AML cell lines revealed a striking and unexpected pattern: the vast majority of TE upregulation occurred through intragenic mechanisms rather than autonomous transcription. Transcriptional readthrough dominated the response, accounting for the largest category of TE activation across all models. This mechanistic bias reflects CDK9’s known role in coordinating not only elongation but also proper 3’ end processing and transcript termination^28^, and therefore these events are unlikely to dominate under other transcriptional perturbations or biological contexts. Consistent with this, applying TEgment to cell cycle-synchronised diploid RPE1 cells, where transcription termination is intact, revealed minimal readthrough-embedded TE expression (Guiducci et al. 2026, submitted). This supports readthrough-driven TE activation as a specific signature of impaired Pol II termination rather than a generic feature of TE transcription. Furthermore, the rapid downregulation of chromatin regulators following CDK9 inhibition likely exacerbates this effect, as epigenetic dysregulation itself has been shown to independently drive widespread transcriptional readthrough and impaired termination^45^. While the next most common modalities were 3’UTR-embedded TEs, here we demonstrated that TEgment is able to identify and quantify previously unannotated TE-derived isoforms (e.g., *LYST*-AluSx, MLT1E1A-*C1orf162*), some of which may produce truncated or altered proteins and affect mRNA stability. Because TEgment not only catalogues but also quantifies these events, it can identify previously unannotated TE-containing isoforms and measure changes in the relative expression of TE-containing versus TE-lacking isoforms of the same gene, enabling isoform-resolved comparison that locus-only approaches cannot access. Contrary to prevailing models linking TE upregulation to autonomous promoter reactivation, standalone TE transcription was a minor fraction of activated loci in CDK9-inhibited AML. TEgment reveals that acute transcriptional perturbation instead triggers TE activation primarily through disrupted co-transcriptional RNA processing.

Moreover, a substantial fraction of TE-containing transcripts in our dataset remained unclassified: transcripts that did not fit cleanly into predefined structural categories. This unclassified fraction potentially highlights the complexity of TE expression patterns that extend beyond current classification schemes and likely includes hybrid or ambiguous transcripts, or multi-TE chimeric structures, incorrectly assembled transcripts, and novel transcripts and isoforms lacking annotation support. Future TEgment development will refine classification logic to capture these edge cases.

Together, these findings argue that “TE expression” should not be interpreted as a single, uniform molecular event: in the setting of acute CDK9 inhibition, increased TE signal can primarily reflect altered transcript architecture and termination rather than activation of autonomous TE promoters.

### Broader applicability and comparison to existing tools

Several computational tools analyse TE-derived transcripts, but each addresses only part of the problem. Quantification-focused tools, including TElocal^15^, Telescope^46^, SQuIRE^47^, and SalmonTE^48^, provide locus- or family-level TE abundance estimates, but treat every TE signal as equivalent, without distinguishing autonomous transcription from passive intronic accumulation, exonization, or readthrough inclusion. Two tools add genomic context to quantification: REdiscoverTE^49^ stratifies repeat-derived reads into exonic, intronic and intergenic pools using reference genome annotation, aggregating them to the subfamily level, while RepExpress^50^ quantifies TEs at the locus level and characterizes each expressed element by the annotated feature it overlaps or, where there is none, by the closest downstream annotated gene. Both are informative and broadly applicable, but they place an element in a genomic context rather than in a transcript: without reconstructing the isoform in which the element resides, an intronic TE carried in unspliced pre-mRNA is not separated from one retained in a mature, spliced transcript. L1EM^51^ does make this distinction, modelling read evidence to separate proper LINE-1 transcription from passive co-transcription, but by design it applies only to L1 retroelements. On the structural side, TEprof3^52^ and LIONS^53^ identify TE-derived alternative promoters, ChimeraTE^54^ and rTEA^55^ resolve TE–gene fusion events, and FREDDIE^56^ identifies TE-containing isoforms without any classification. These tools are powerful within their respective niches, but each was designed for a specific structural event, and none offers a unified framework in which the full spectrum of TE expression modes can be quantified and compared within the same sample. TEgment consolidates these capabilities with locus-resolved quantification, structural annotation across all modalities, and statistical differential expression into a single reproducible workflow.

Because TEgment reconstructs transcript isoforms *de novo*, it reports only loci with structural support and recovers TE-containing isoforms that are absent from the reference annotation (Figure 5). The cost is that it is more demanding of input data (see Limitations and future directions) than assembly-free tools such as REdiscoverTE and RepExpress, which remain preferable for shallow, single-end or fragmented libraries and for species lacking GENCODE annotation, and that genuine TE expression falling outside assembled transcripts is not reported. We therefore regard the two strategies as complementary.

The computational framework established here has implications that extend well beyond CDK9 inhibition. TEgment is designed to be broadly applicable across any biological context in which TE expression is of interest, from development and aging to viral mimicry and immunotherapy response. Importantly, the structural classification approach will yield different predominant modalities in different contexts: readthrough will be more prominent when transcription termination is impaired; UTR and lncRNA embedding will dominate when specific host gene programs are activated; autonomous standalone transcription may become more prominent in contexts of global DNA demethylation or SETDB1 loss. TEgment provides the quantitative framework to make these distinctions systematically and comparably across studies.

Moreover, TEgment supports RNA-seq library types suitable for short-read transcriptome assembly, including steady-state RNA-seq and nascent RNA-seq. It accommodates both single-end and paired-end reads (paired-end recommended), with flexible read lengths. TEgment handles multi-mapping reads via expectation-maximization model, and users can alternatively restrict quantification to uniquely mapped reads to obtain conservative calls. This flexibility makes TEgment suitable for diverse transcriptomic datasets and experimental designs.

### Limitations and future directions

While TEgment represents a significant advance in TE transcriptome analysis, some limitations warrant consideration. First, the reference-guided transcriptome assembly step relies on StringTie3 and therefore benefits from stranded, 150 bp paired-end libraries with sufficient depth; sparse or single-end data will yield less reliable isoform reconstruction. Furthermore, the pipeline depends on reference annotation to classify TE–gene relationships and may miss novel TE-containing transcripts, especially poorly annotated lncRNAs and TE insertions that are absent from the reference genome, including germline-polymorphic and somatically acquired, cell-line-specific insertions. Relatedly, TEgment does not currently annotate L1-mediated 3′ transduction events as a distinct category; such transcripts are quantified but classified according to their downstream flanking context.

Second, the current implementation of TEgment does not detect TE–TE junction events, chimeric transcripts formed by the fusion of multiple TE copies or different TE families. Such events, which may arise from trans-splicing or genomic rearrangements represent an additional layer of TE-mediated transcriptomic diversity that we are implementing in the future version of TEgment. Additionally, classifications such as TE-initiation, TE-termination, exonized, and exonic currently work only for protein-coding genes, not for non-coding RNAs. The classification logic is also deterministic, so transcripts that straddle category boundaries (for example, a TE that is simultaneously exonized and acting as a promoter for another isoform) are forced into a single bin; a probabilistic, multi-label scheme would recover these ambiguous events.

Third, short-read sequencing limits the resolution of full-length transcript structures. Moreover, locus-level quantification may not be reliable for the youngest and least divergent TEs. For these elements, family-aggregation provides the more reliable resolution, and long-read RNA sequencing (Oxford Nanopore, PacBio) will be essential for definitive isoform characterization and multi-mapping resolution in the transcriptome assembly step and will be integrated into future versions of TEgment. Fourth, the TEgment classification module depends heavily on high-quality genomic annotation. To ensure accurate results, TEgment currently supports only GENCODE annotation data, which restricts its application exclusively to human and mouse genomes. In future updates, we plan to integrate support for RefSeq and Ensembl annotations to extend the tool’s utility to non-model organisms.

## Lead contact

Further information and requests for resources should be directed to and will be fulfilled by the Lead Contact, Özgen Deniz.

## Material availability

This study does not generate new unique reagents.

## Acknowledgments

We thank Deniz lab members for constructive feedback and Paul Hurd, Miguel Branco, Lovorka Stojic and John Gribben for their comments on the manuscript. All computational works were performed on QMUL’s high-performance computing cluster, Apocrita. Ö.D. and her group receives funds from a Cancer Research UK Career Development Fellowship (RCCCDF-Nov21\100002) and UK Research and Innovation (UKRI)’s Horizon Europe Guarantee scheme (EP/Y030338/1). All figures were created with Inkscape and Biorender.

## Author contributions

Conceptualization, D.P., and Ö.D.; software, D.P.; formal analysis, D.P.; investigation: A.M, D.P., H.T, E.L., and Ö.D.; data curation, D.P, and Ö.D.; visualization, D.P.; writing – original draft D.P, and Ö.D.; writing – review&editing, all authors; resources, Ö.D; funding acquisition, Ö.D.; supervision: Ö.D.; project administration, Ö.D.

## Declaration of generative AI and AI-assisted technologies in the manuscript preparation process

During the preparation of this work, the authors used Anthropic Claude Opus to improve language clarity and assist with data analysis scripts. The authors reviewed and edited the output as needed and take full responsibility for the content of the published article.

## Supplemental information

Document S1. Figures S1–S8

Table S1. Gene expression dataset sample accession IDs and publicly available CDK9 inhibitor-treated RNA-seq data, related to Figure 1.

Table S2. Gene composition in MONO-MAC-6 time course clusters, related to Figure 3.

Table S3. TE composition in MONO-MAC-6 time course clusters, related to Figure 3.

