## Supplementary document for "TEgment dissects transposable element reactivation upon CDK9 inhibition in acute myeloid leukemia"

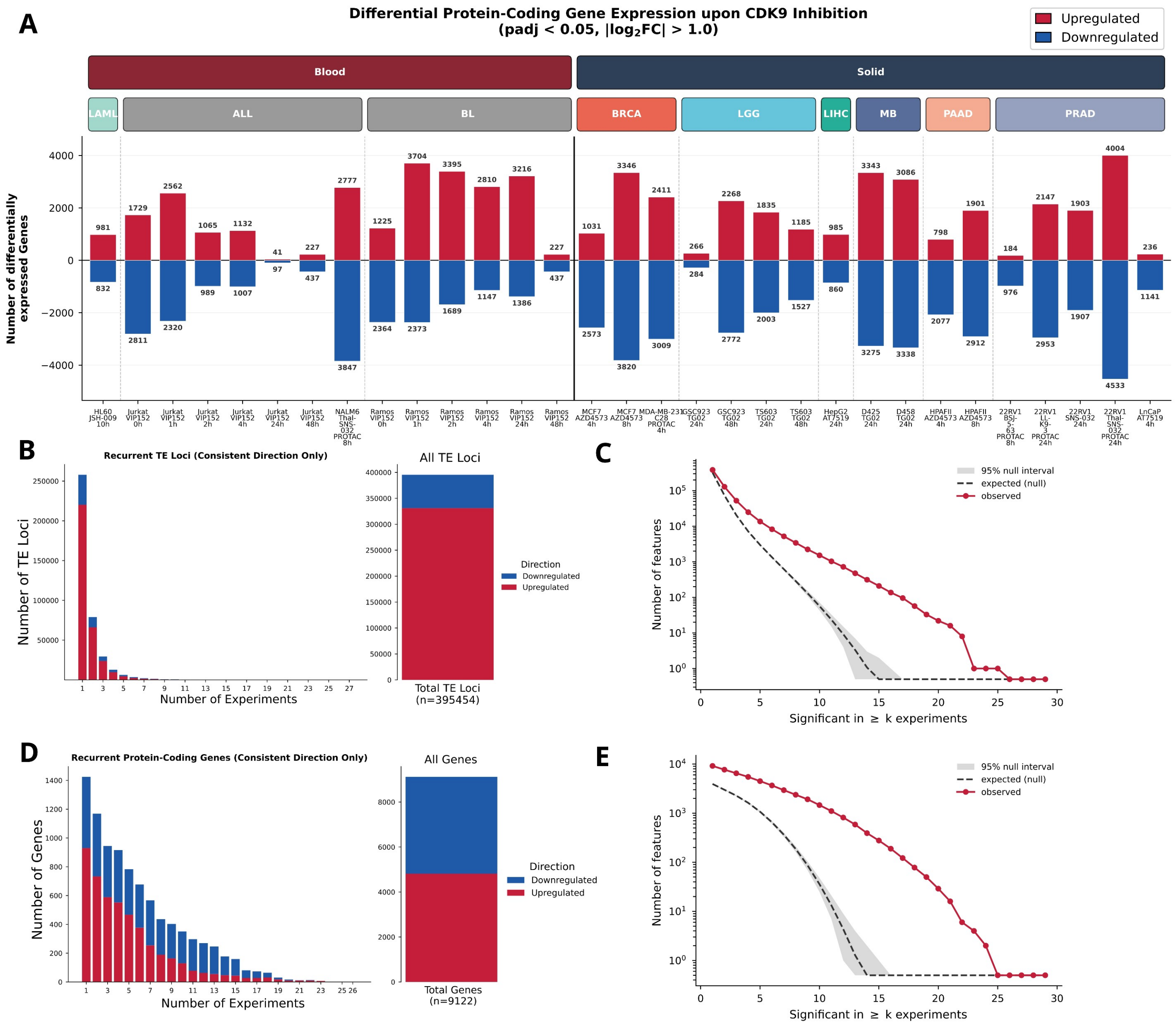


**Supplementary Figure 1. Global transcriptomic alterations upon CDK9 inhibition, related to Figure 1. (A)** Number of differentially expressed protein-coding genes following CDK9 inhibition. Data are stratified by major tumor classification (Blood vs. Solid) and specific cancer lineages (LAML, ALL, BL, BRCA, LGG, LIHC, MB, PAAD, PRAD). Red bars represent upregulated genes, while blue bars indicate downregulated genes. Differential gene expression was determined using DESeq2 (padj < 0.05, |log$_2$ Fold Change| > 1.0). **(B)** Recurrence analysis of individual transposable element (TE) loci. The histogram displays the number of distinct TE loci (y-axis) showing consistent, unidirectional expression changes across multiple independent experiments (x-axis). The adjacent stacked bar chart summarizes the total pool of differentially expressed TE loci, categorized by overall directionality (red = upregulated, blue = downregulated). **(C)** Permutation testing of TE locus recurrence. Observed number of TE loci significant in the same direction in ≥k experiments (red) compared with the mean (dashed line) and 95% interval (grey) of 10,000 expression-matched permutations. **(D)** Recurrence analysis of differentially expressed protein-coding genes, plotted as in (B). The histogram illustrates the frequency of genes (y-axis) exhibiting consistent, unidirectional dysregulation across multiple independent experiments (x-axis); the adjacent stacked bar chart summarizes the total pool of these genes by directionality. **(E)** Permutation testing of protein-coding gene recurrence, plotted as in (C).

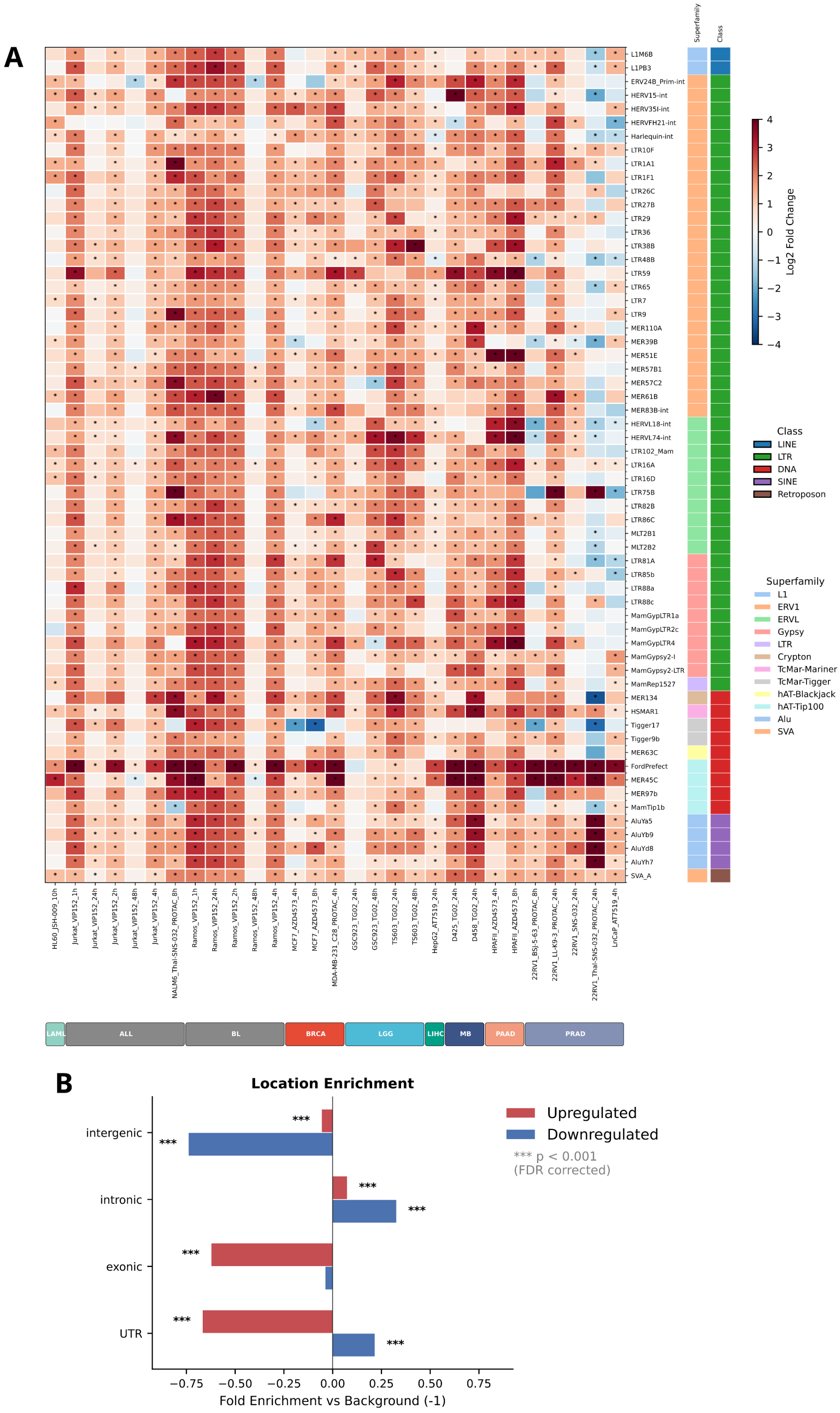


**Supplementary Figure 2. Family level expression dynamics and genomic location enrichment of TEs upon CDK9 inhibition, related to Figure 1.**

**(A)** Differential expression profiles of highly recurrent TE families across diverse cancer models. Columns represent individual experimental conditions, categorized by cancer lineage. Rows correspond to specific TE families. The central color scale signifies the log2 fold change, with red indicating upregulation and blue indicating downregulation. Cells marked with an asterisk denote statistically significant differential expression (adjusted p-value $< 0.05$ and $|\log_2 \text{fold change}| > 1.0$). Color-coded row annotations on the right map each TE family to its corresponding superfamily and major TE class. **(B)** Genomic location enrichment analysis of consistently upregulated (red) and downregulated (blue) TE loci relative to the expected distribution of all RepeatMasker-annotated TEs (background). The x-axis displays the fold enrichment, where positive values indicate enrichment and negative values indicate depletion within the specified genomic regions (Intergenic, Intronic, exonic, and UTR). Statistical significance was assessed using Fisher's exact test followed by Benjamini-Hochberg FDR correction (*** $p < 0.001$).

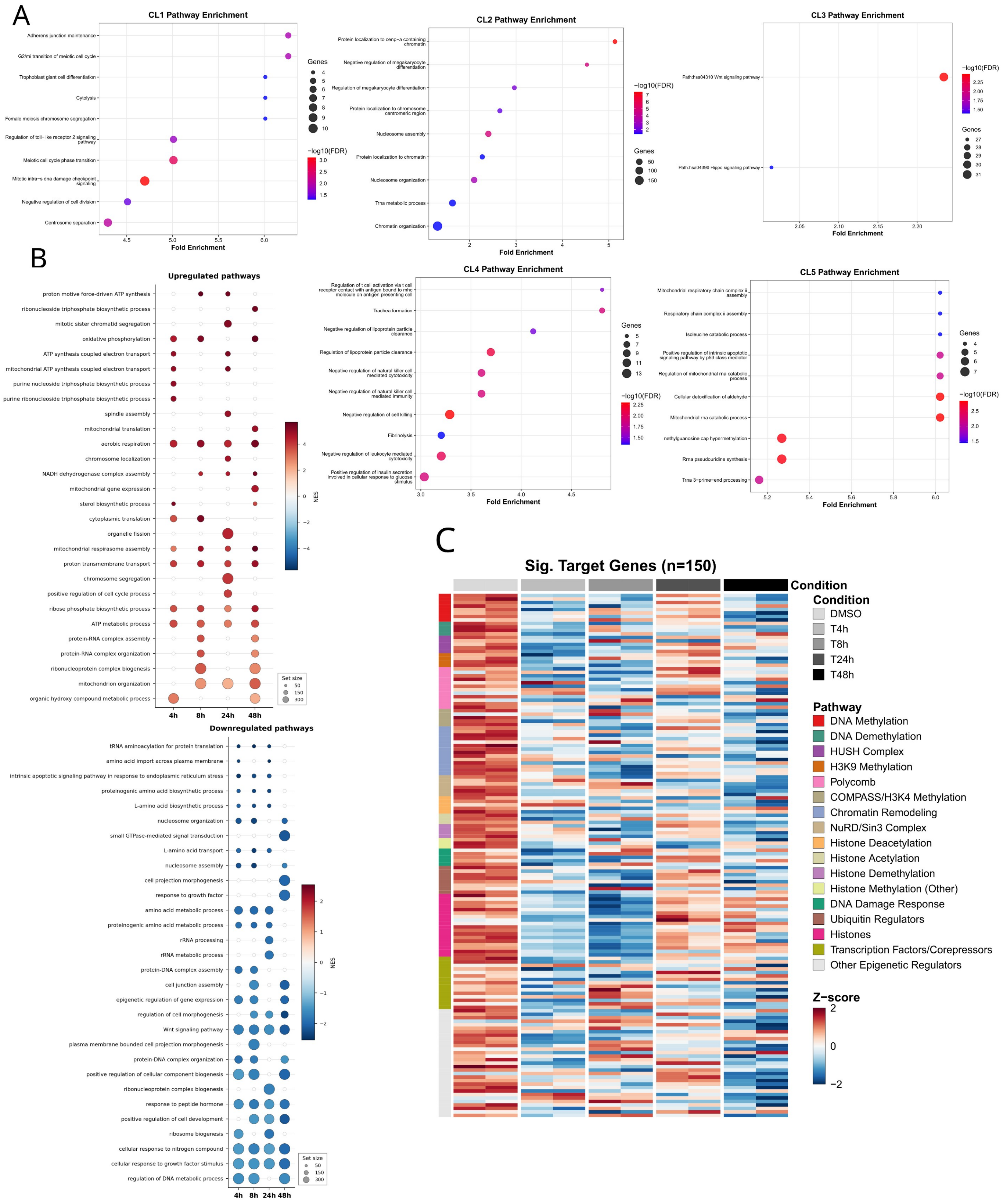


### Supplementary Figure 3. Pathway enrichment and dynamic regulation of epigenetic modifiers following CDK9 inhibition, related to Figure 3

**(A)** Pathway enrichment analysis for the five distinct transcriptional clusters (CL1–CL5) identified during the CDK9 inhibition time course. Bubble plots display the top enriched Gene Ontology (GO) biological processes for each cluster. The x-axis represents Fold Enrichment. Bubble size corresponds to the number of significant genes overlapping with the respective pathway (Genes). Bubble color indicates statistical significance based on $-\log_{10}$(FDR), utilizing a gradient from blue (lower significance) to red (higher significance). **(B)** GSEA of transcriptional changes across the AZD4573 treatment time course relative to the DMSO control. The dot plots summarize the top upregulated (top panel) and downregulated (bottom panel) biological pathways over time. Dot size reflects the total number of genes in the gene set (Set size). Dot color represents the NES, with red gradients indicating positive enrichment (upregulated pathways) and blue gradients indicating negative enrichment (downregulated pathways). **(C)** Temporal expression dynamics of significantly differentially expressed epigenetic regulators (Sig. Target Genes, LRT FDR $< 0.05$). Columns represent individual biological replicates across the treatment conditions (DMSO, T4h, T8h, T24h, T48h). Rows represent individual genes. Expression values are variance-stabilized and row-scaled (Z-score). The left annotation bar categorizes each gene into specific epigenetic regulatory pathways and complexes, as detailed in the accompanying legend key.

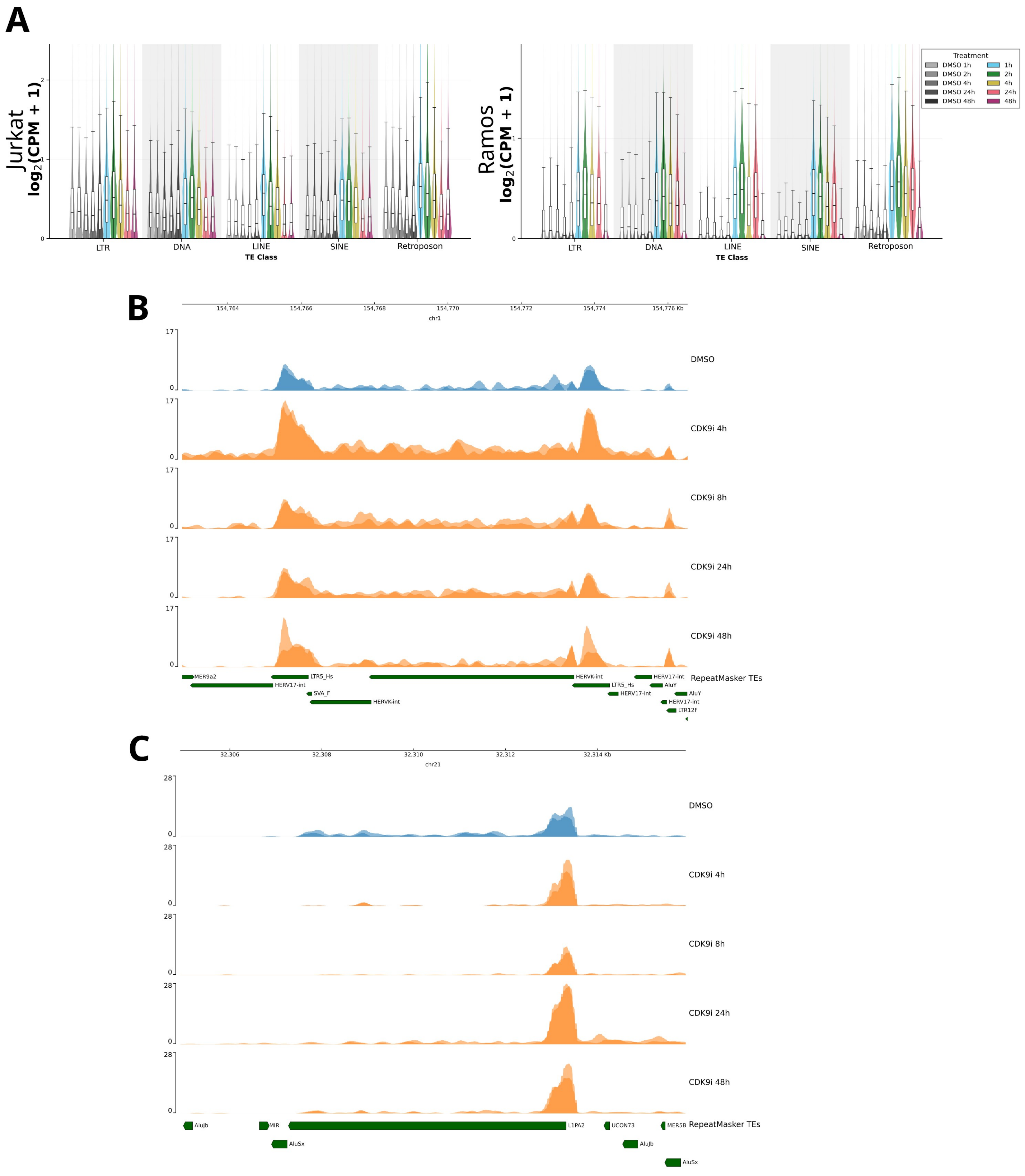


**Supplementary Figure 4. Dynamics of transposable element expression following CDK9 inhibition, related to Figure 3**

**(A)** Distribution of $\log_2$(CPM + 1) expression for expressed transposable element (TE) loci in Jurkat (left) and Ramos (right) cell lines treated with a CDK9 inhibitor

VIP152. TEs are categorized by class. Expression is shown across a time course. Box plots within the violins show the median and IQR. **(B-C)** Representative genome tracks showing induction of TEs following CDK9 inhibition in MONO-MAC-6 cells. Tracks show normalized RNA-seq read coverage. Two biological replicates are overlaid for each condition: DMSO control (blue) and CDK9 inhibitor treatment (orange). The bottom track (green) displays RepeatMasker annotations, highlighting individual TE loci within the corresponding genomic regions.

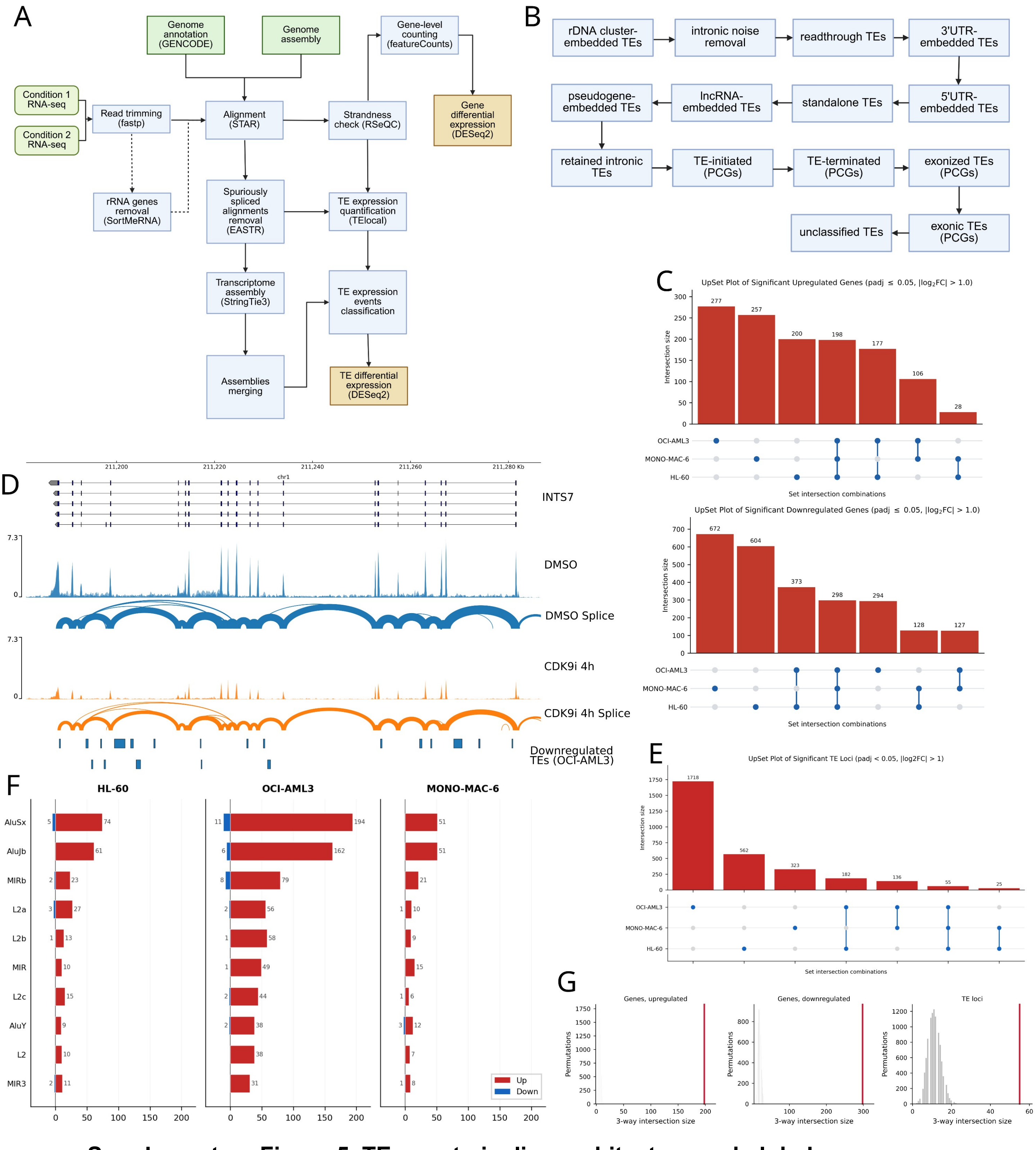


**Supplementary Figure 5. TEgment pipeline architecture and global transcriptomic dynamics across AML models, related to figure 4.**

**(A)** Schematic overview of the TEgment modular analysis pipeline, detailing data processing from raw RNA-seq reads to gene and locus level TE differential expression. **(B)** Flowchart outlining the hierarchical, rule-based classification module used to map the structural modalities of TE expression events. PCG: protein-coding genes. **(C)** Overlap of significantly upregulated (top) and downregulated (bottom) protein-coding genes across OCI-AML3, MONO-MAC-6 (MM6), and HL-60 cell lines following 4h CDK9 inhibition. **(D)** Representative genome browser track demonstrating pervasive transcriptomic noise and passive intronic background signal in the gene *INTS7* in OCI-AML3 following CDK9 inhibition. Tracks show normalized RNA-seq read coverage. Two biological replicates are overlaid for each condition: DMSO control (blue) and CDK9 inhibitor treatment (orange). The bottom track (blue) displays significantly downregulated TEs. **(E)** Intersection of significantly dysregulated TE loci across the three AML cell lines ($padj < 0.05$, $|log2FC| > 1$). **(F)** Absolute frequency of significantly upregulated (red) and downregulated (blue) TEs, categorized by the most consistently enriched TE families across the three AML models. (G) Null distributions of the three-way intersection between HL-60, MONO-MAC-6 and OCI-AML3 for upregulated genes (left), downregulated genes (middle) and dysregulated TE loci (right), from 10,000 label permutations; red line, observed value.

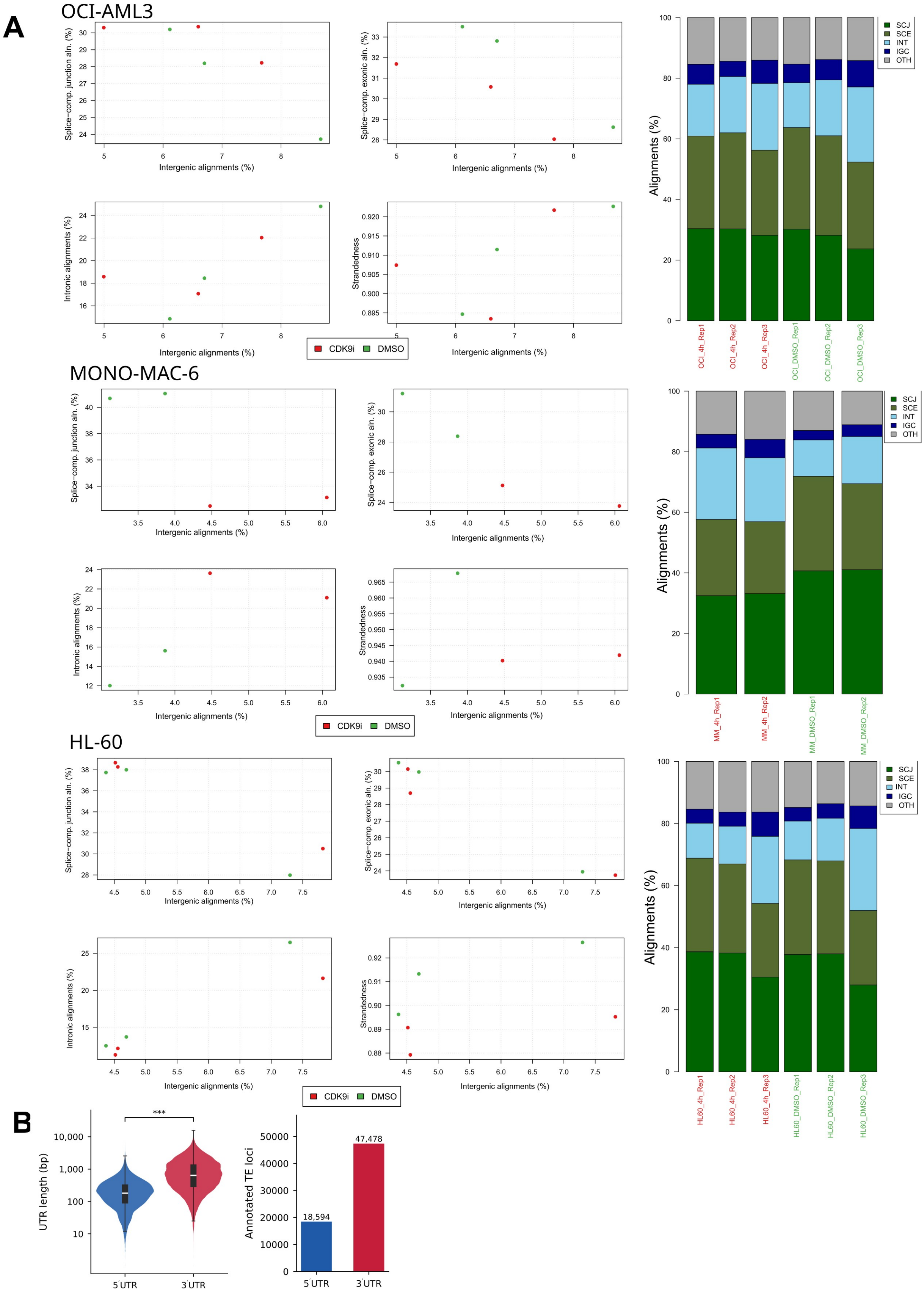


**Supplementary Figure 6. RNA-seq library diagnostics and UTR compartment size, related to Figure 4.**

(A) gDNAx diagnostics for OCI-AML3, MONO-MAC-6, and HL-60 CDK9i and DMSO libraries. Left: intergenic alignment rate versus splice-compatible junction, splice-compatible exonic, and intronic alignment rates, and library strandedness. Right: per-sample alignment composition. SCJ, splice-compatible junction; SCE, splice-compatible exonic; INT, intronic; IGC, intergenic; OTH, other. (B) UTR length (left) and number of overlapping RepeatMasker-annotated TE loci (right) for 5′ and 3′UTRs of protein-coding transcripts (GENCODE v49). ***$P < 0.001$, Mann–Whitney U test.

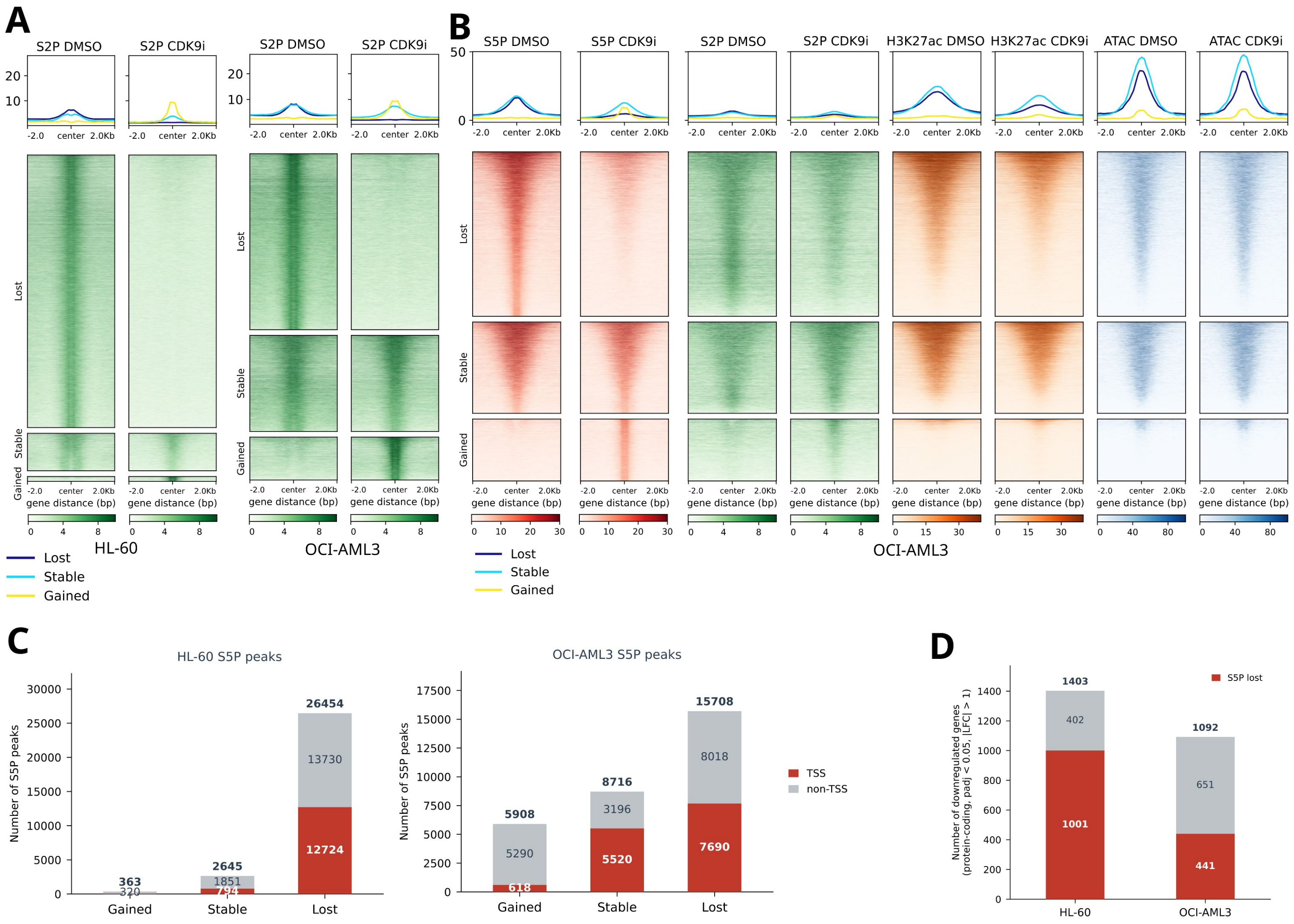


**Supplementary Figure 7. Chromatin and transcriptional dynamics following CDK9 inhibition in AML cell lines, related to figure 6.**

**(A)** Pol II-S2P signal intensity in HL-60 and OCI-AML3 cells treated with DMSO or a CDK9 inhibitor (CDK9i). Signal is centered around S2P peak regions and stratified into “Lost”, “Stable”, and “Gained” categories. **(B)** Distribution of Pol II-S2P, Pol II-S5P, H3K27ac, and ATAC-seq signals in DMSO and CDK9i-treated OCI-AML3 cells. The data is centered around Pol II-S5P peak sites and categorized by S5P status (Lost, Stable, or Gained). **(C)** Total number of S5P peaks classified as Gained, Stable, or Lost following CDK9i treatment in HL-60 (left) and OCI-AML3 (right) cells. Peaks are further annotated by their genomic location, distinguishing between those at TSS (red) and non-TSS regions (gray). **(D)** Number of significantly downregulated protein-coding genes (adjusted p-value < 0.05, |log2FoldChange| > 1) following CDK9i treatment in HL-60 and OCI-AML3 cell lines. The bars illustrate the proportion of these downregulated genes that exhibit a corresponding loss of S5P signal (red) versus those without S5P loss (gray).

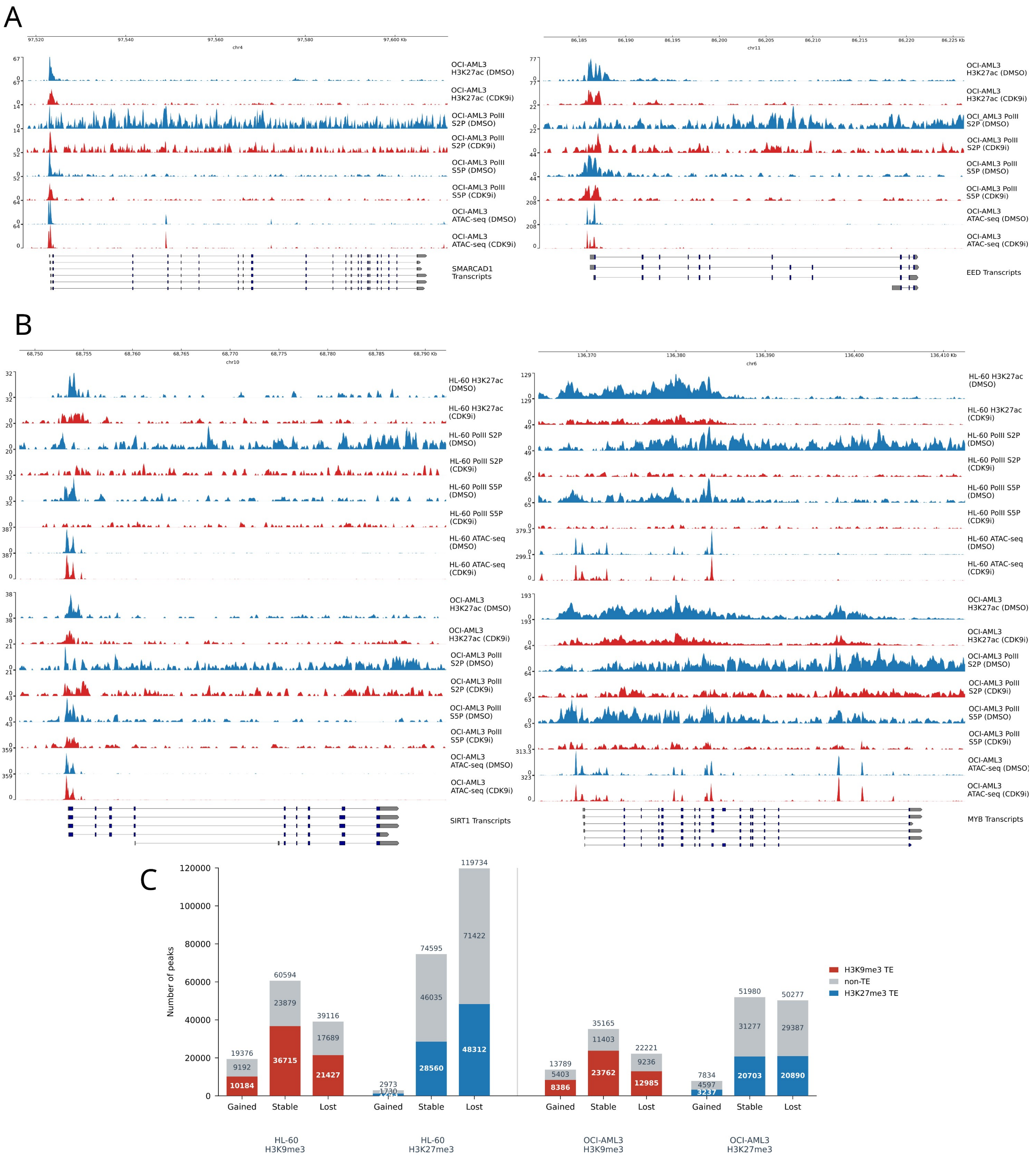


**Supplementary Figure 8. Epigenomic and transcriptional changes at representative gene loci and genome-wide heterochromatin dynamics following CDK9 inhibition, related to figure 6.**

**(A)** Signal distribution of H3K27ac, Pol II-S2P, Pol II-S5P, and ATAC-seq signals in OCI-AML3 cells. Cells were treated with either DMSO (control, blue tracks) or a CDK9 inhibitor (CDK9i, red tracks). The panels show representative genomic loci for the *SMARCAD1* (left) and *EED* (right) genes. **(B)** H3K27ac, Pol II-S2P, Pol II-S5P, and ATAC-seq signals in both HL-60 (top sets of tracks) and OCI-AML3 (bottom sets of tracks) cell lines. Signal is shown for both DMSO (blue) and CDK9i (red) treatment conditions at the *SIRT1* (left) and *MYB* (right) gene loci. **(C)** Total number of H3K9me3 and H3K27me3 heterochromatin peaks classified as Gained, Stable, or Lost after CDK9 inhibition in HL-60 and OCI-AML3 cells. The bars indicate the proportion of peaks overlapping with TE versus non-TE genomic regions. Red segments represent H3K9me3 TEs, blue segments represent H3K27me3 TEs, and gray segments represent non-TE regions.
